# Higher frequency of homologous chromosome pairing in adult endothelial cells as compared to neonatal endothelial cells

**DOI:** 10.1101/2025.03.15.643486

**Authors:** Jemery Morales, Omar Akhtar, Gabriel Quintero Plancarte, Lisa Hua

**Affiliations:** Biology Department, Sonoma State University, Rohnert Park, CA 94928

**Keywords:** Antipairing, homologous chromosomes, human aortic endothelial cells, human umbilical vein endothelial cells, mitosis

## Abstract

We previously demonstrated that homologous chromosomes are spatially segregated, or antipaired, in neonatal human endothelial cells at metaphase/anaphase, which may help prevent abnormal recombination. However, it is unclear if this antipairing persists in adult endothelial cells. To test whether the antipairing, or one homolog per nuclear hemisphere motif, is conserved in adult endothelial cells, we examined human aortic endothelial cells at metaphase. Using ImmunoFISH and high-resolution confocal microscopy to visualize the chromosomes and centrosomes, we found that small homologous chromosomes 13, 15, 17, 19, 21, 22, and the sex chromosomes, XY, exhibit a loss of spatial segregation in human adult aortic endothelial cells. In addition, fewer adult endothelial cells showed segregation for the larger chromosomes 1, 4, and XX, as compared to neonatal endothelial cells. Notably, we observed a higher frequency of abnormal pairing in both small and large chromosomes in adult aortic endothelial cells as compared to neonatal umbilical vein endothelial cells. These findings suggest that mechanisms governing chromosome antipairing may decline with aortic endothelial cell age.

**Summary Statement:** A higher frequency of homologous chromosome pairing is observed in human adult aortic endothelial cells as compared to neonatal umbilical vein endothelial cells.

## Introduction

Chromosomal organization is important for endothelial cell (EC) function, particularly during mitosis, when a cell divides to produce two identical daughter cells and maintain the correct chromosome number across cell divisions (Hua et al., 2022; Srinivasan et al., 2022). Non-random chromosome organization may play a role in maintaining genomic stability (Cai, Casas, Quintero Plancarte et al., 2025; Cremer and Cremer, 2001; Cremer et al., 2020; Hua and Mikawa, 2018b; Hua et al., 2022; Koeman et al., 2008; Meaburn et al., 2007; Misteli and Soutoglou, 2009). Research from our lab has shown that homologous chromosomes in neonatal human umbilical vein endothelial cells (HUVECs) are arranged on opposite sides of the centrosome axis during metaphase/early anaphase, resulting in a haploid set on either side (Cai, Casas, Quintero Plancarte et al., 2025; Hua and Mikawa, 2018b). This haploid set organization, or antipairing pattern, may minimize homologous chromosome pairing, indicating its importance in early development. However, it is unknown whether this antipairing pattern persists throughout adulthood.

Segregation of haploid genomes has also been observed in other model systems including the mouse, bovine, rabbit, fruit fly, and nematode (Gondos and Bhiraleus, 1970; Hua et al., 2022; Joyce et al., 2012; Joyce et al., 2016; Keneklis and Odartchenko, 1974; Mayer et al., 2000a,b; Maddox et al., 2005; Odartchenko and Keneklis, 1973; Reichmann et al., 2018; Schneider et al., 2021; Velez-Aguilera et al., 2020). This haploid set organization defines the earliest stages of development, with the observation of parental genome separation in mammalian models (Mayer et al., 2000a,b; Reichmann et al., 2018; Schneider et al., 2021). For example, in mouse and bovine cells, dual mitotic spindles at fertilization separate the two parental sets (Reichmann et al., 2018; Schneider et al., 2021). In addition, 5-bromodeoxyuridine (BrdU)-labeled sperm show paternal genome separation in mouse zygotes which persist until the four-cell stage (Mayer et al., 2000b). Electron microscopy of fertilized rabbit ova shows intact parental pronuclei adjacent to each other until approximately 21 hours after mating (Gondos and Bhiraleus, 1970). After fertilization in the fruit fly, gonomeric cell divisions show the maternal and paternal genomes in apposition and structurally separate (Callaini and Riparbelli, 1996; Loppin et al., 2015). An evolutionary persistence may reflect selective pressure to limit harmful recombination and promote nuclear compartmentalization. In this study, we observe a degradation of the haploid set pattern postnatally.

Antipairing may be an important process in the fruit fly, as it may be inhibited or enhanced by different regulators, suggesting there are mechanisms driving the antipairing of homologous chromosomes in different animal models (Joyce et al., 2012; Joyce et al., 2016). For example, in the early fruit fly embryo, a high-throughput FISH-based screen identifies genes involved in promoting or antagonizing somatic homolog pairing (Joyce et al., 2012). This finding suggests that there may be active antipairing mechanisms to prevent the pairing of chromosomes in mammals. Taken together, these findings support the concept that antipairing is a biologically significant nuclear architecture strategy to maintain overall genomic stability.

Changes in chromatin structure further alter gene expression and increase susceptibility to DNA damage, with aging associated with higher levels of damage due to oxidative stress (Burgess et al., 2012; Wood and Helfand, 2013; Wood et al., 2010; Zhang et al., 2021). Notably, the spatial arrangement of chromosomes within the cell may become disorganized in aged ECs, which can impact gene expression and overall cellular function (Ghosh and Meyer, 2021; Heard and Bickmore, 2007; Hua et al., 2022; Koeman et al., 2008; Lima et al., 2022; Misteli, 2007). Aging disrupts nuclear organization through the loss of heterochromatin and nuclear lamina integrity, causing gene-rich regions to shift away from transcriptionally active domains and altering gene expression patterns (Chandra and Kirschner, 2016; Emerson and Lee, 2023; López-Otín et al., 2013).

The loss of antipairing may increase the chances of homologous chromosome pairing, leading to mitotic recombination and potential genomic instability. This loss has been correlated with renal cancer and altered gene expression (Hua and Mikawa, 2018b; Koeman et al., 2008). This finding suggests that abnormal somatic pairing and the loss of antipairing could be related to various human pathologies (Atkin and Jackson, 1996; Faruqi et al., 1994; Hua and Mikawa, 2018b; Hua et al., 2022; Koeman et al., 2008; Lewis et al., 1993). Despite this, there has been no thorough investigation into homologous chromosome organization in aged ECs, and a comprehensive analysis of this organization during mitosis is still lacking.

In this study, we employed 3D imaging to examine the organization of homologous chromosomes in adult human aortic endothelial cells (HAECs) sourced from individuals, both female and male, in the age range of 21 years to 68 years, and cultured at low passage numbers. Using immunofluorescence and DNA fluorescence *in situ* hybridization (ImmunoFISH) and high-resolution confocal microscopy, we analyzed the positions of homologous chromosomes to determine whether the spatial segregation of homologous chromosomes, or antipairing, persists in adult aortic ECs. Our findings reveal that mitotic antipairing is lost in adult aortic ECs. Small chromosomes 13, 15, 17, 19, 21, 22, and XY lost their spatial segregation in HAECs. In addition, fewer HAECs showed segregation as compared to HUVECs for the larger chromosomes 1, 4, and XX. This suggests a gradual decline in the fidelity of spatial segregation of homologous and sex chromosomes in adult ECs. Both small and large chromosomes exhibited a higher frequency of abnormal homologous pairing in adult HAECs compared to neonatal HUVECs. The smaller chromosomes, including the acrocentric ones, showed a greater frequency of abnormal pairing than larger chromosomes in adult HAECs. There are at least two potential causes for this higher pairing frequency and the loss of antipairing: one would be an age-dependent change, and the other would be an aorta-specific mechanism to enhance the loss of antipairing. Overall, our data indicate that the mechanisms governing chromosome antipairing organization may decline, or lose fidelity, in aged aortic ECs.

## Results

### Spatial segregation was maintained for small chromosomes 17 and 19 in neonatal human umbilical vein endothelial cells (HUVECs) at metaphase

Previously, it was found that a pair of homologous chromosomes resides on opposite sides of the centrosome, or nuclear division axis, in human neonatal cells at metaphase/early anaphase (Hua and Mikawa, 2018b). This one homolog on either side of the centrosome axis, or antipairing, organization was conserved for all chromosomes in the human karyotype of HUVECs at early anaphase (Hua and Mikawa, 2018b). To confirm that antipairing is present at metaphase, individual chromosomes were visualized in HUVECs (Fig. 1A). HUVECs are polarized cells with an apical and basal domain (Muller and Gimbrone, 1986), which aided in the establishment of an axial coordinate system with a chromosome paint approach (Hua and Mikawa, 2018b). In addition, ImmunoFISH assays were performed to simultaneously visualize the centrosome axis (Fig. S1).

**Fig. 1.**
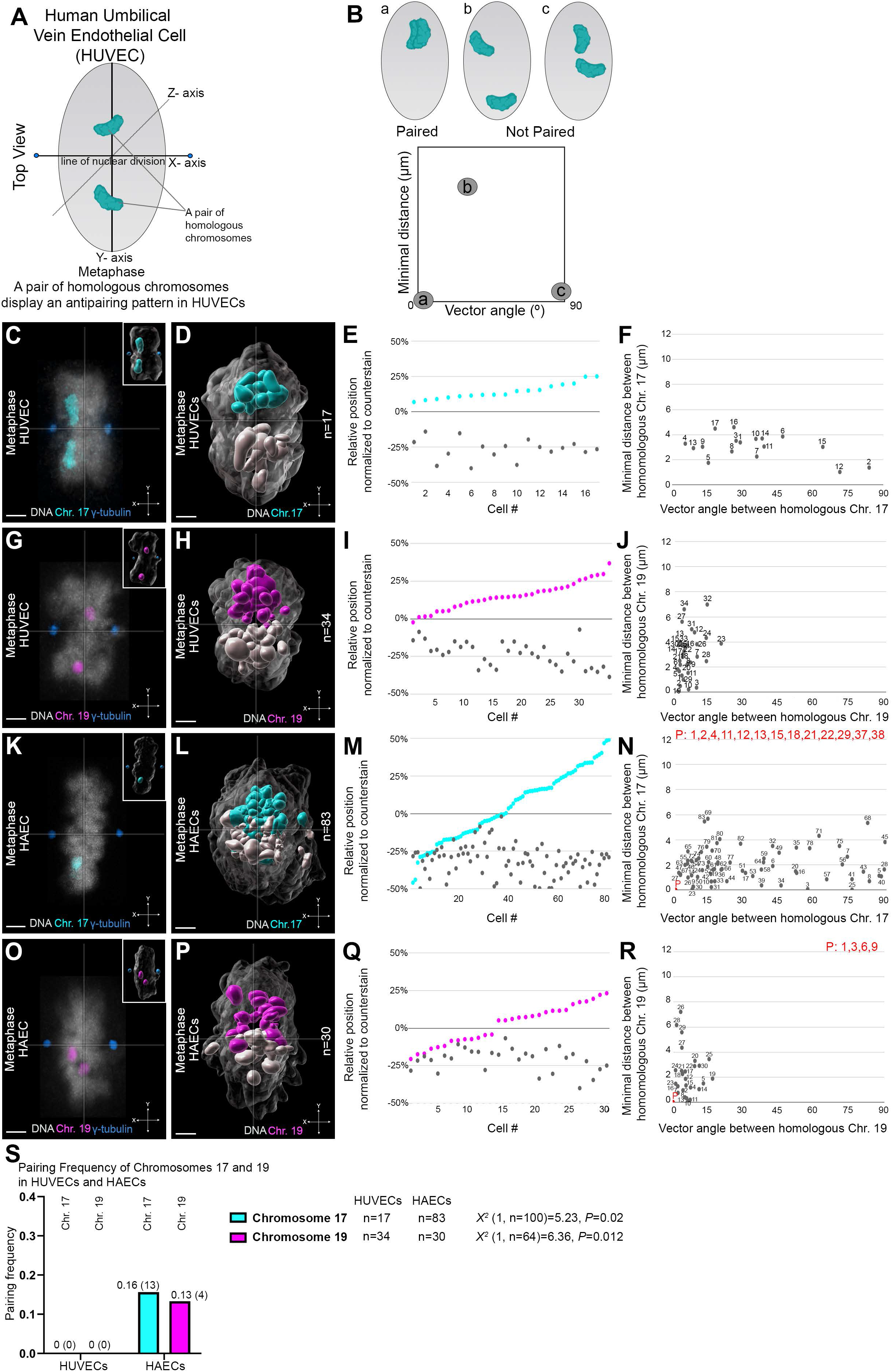
Homologous chromosome segregation was lost in small chromosomes 17 and 19 in adult HAECs. **(A)** Schematic showing a pair of homologous chromosomes localizing on opposite sides of the nuclear hemisphere for both autosomal, and sex chromosomes in neonatal human umbilical vein endothelial cells (HUVECs) (Hua and Mikawa, 2018b). **(B)** Pairing is defined as the distance between two homologous chromosomes at 0 µm and vector angle at 0°. Different scenarios of paired (a) vs not paired (b, c) are shown. **(C)** Top view of optical sections of a HUVEC at metaphase stained for chromosomes 17 (cyan), γ-tubulin (blue), and counterstained for DNA (grey). Inset: 3D reconstruction of the optical sections. **(D)** A 3D overlay of chromosome 17 (cyan/grey) of multiple HUVECs at metaphase, where the homolog closest to the x-axis (y=0) is colored (cyan), while its respective partner is in grey (segregation in n=17/17 cells). **(E)** Relative positions for each homolog of chromosome 17 (cyan/grey circles) when mapped to an axial coordinate system. **(F)** The distance and angular orientation of homologous chromosome 17 pairs with an average distance of 3.03±0.99 µm s.d. and angular orientation of 34.5°±21.9° s.d. *Note: Each numbered cell corresponds to the same cell number in the relative position normalized to the DNA counterstain.* **(G-J)** As in C-F, but of chromosome 19 (segregation in n=33/34 cells). The distance and angular orientation of chromosome 19 homologs with an average distance of 2.98±1.70 µm s.d. and angular orientation of 5.61°±4.27° s.d. **(K-N)** As in C-F, but of adult human aortic endothelial cells (HAECs) (n=4 patients, segregation in n=44/83 cells) with an average distance of 1.68±1.40 µm s.d. and angular orientation of 27.2°±27.5° s.d. *Note: Red P indicates pairing and the following numbers are cells in which pairing has been identified*. **(O-R)** As in G-J, but of HAECs. The distance and angular orientation of chromosome 19 homologs (n=4 patients, segregation in n=17/30 cells) with an average distance of 1.99±1.89 µm s.d. and angular orientation of 5.43°±4.53° s.d. **(S)** Pairing frequency graph showing the pairing of chromosomes 17 (cyan) *X^2^* (1, n=100)=5.23, *P*=0.02 and 19 (magenta) *X^2^* (1, n=64)=6.36, *P*=0.012 of HUVECs and HAECs. Pairing data were analyzed using binomial logistic regression. Scale bar: 2 µm.

The nuclear hemispheres were previously defined by Hua and Mikawa (2018b) as the following: the x-axis is the line that crosses the centrosomes, the z-axis is the laser line, and the y-axis is perpendicular to both the x- and z-axes at metaphase/anaphase (Fig. 1A; Fig. S2A’-A”’). To evaluate the frequency of homologous pairing, we measured the minimal distance between each homologous pair as previously described (Hua and Mikawa, 2018b) (Fig. 1B). Pairing is defined as having a 0 µm distance and a 0° vector angle, determined in a 3D space, between a pair of homologous chromosomes (Fig. 1B). Thus, the spatial proximity of 0 µm distance and 0° vector angle is interpreted as direct spatial juxtaposition or contact along the axis of each homolog in our analysis (Fig. 1B). P and q arm orientation was not used to determine angular orientation of homologs due to the resolution of the chromosome paint probes that bind to sequences across the chromosome without distinction of p and q regions.

Our previous analysis of angular orientation was completed using a 180° range (Cai, Casas, Quintero Plancarte et al., 2025; Hua and Mikawa, 2018b). Angular orientation is determined by the angle between the 3D vectors along each homolog. Each vector starts at the point of origin of the centromere, and runs through the chromosome arm via the longest ellipsoid axis (Hua and Mikawa, 2018b). The vector angle was calculated by taking the inverse cosine of the dot product of two 3D vectors divided by the product of their magnitudes. At anaphase, spindle microtubules attach to kinetochores that surround centromeres of individual homologs that pull sister chromatids apart (Magidson et al., 2011; Walczak et al., 2010). This leads to chromosome arms trailing behind the centromeres (Magidson et al., 2011; Walczak et al., 2010).

There is less constraint of chromosome arms at metaphase and more dynamic movement (Cai, Casas, Quintero Plancarte et al., 2025; Magidson et al., 2011). At metaphase, the centromere position was less certain relative to the rest of the chromosome arm as centromeres/kinetochores oscillate parallel to the centrosome axis (Cai, Casas, Quintero Plancarte et al., 2025; Magidson et al., 2011). This stage difference makes it harder to identify the centromere position in fixed samples at metaphase, and does not allow us to assign directionality of vectors at the 180° range. Therefore, we limited our analysis to a single quadrant (90° range).

To determine whether the homologous chromosomes were segregated at metaphase, we performed ImmunoFISH to combine immunolabeling using a γ-tubulin antibody to visualize the centrosomes (O’Toole et al., 2012), DNA FISH/Whole Chromosome Paint (WCP) to visualize chromosomes, and a DNA counterstain (Fig. 1C,G,K,O). For WCP experiments, the centrosomes were not able to be visualized. Therefore, a DNA-derived axis perpendicular to the longest axis of the 3D metaphase DNA plate was used to determine the centrosome axis. Immunofluorescence staining for γ-tubulin, along with a DNA counterstain, revealed that the centrosome axis and DNA-derived axis had a difference of 0.5 µM (Fig. S1). Consequently, individual chromosome positions within ± 0.5 µM of the DNA center of volume, referred to as the “boundary region”, were excluded from the analysis. For instance, out of 66 total cells, 6 cells exhibited homologous chromosome 1 in this boundary region, rendering the data uninterpretable (n=6/66 cells for chromosome 1).

We then determined if a pair of homologous chromosomes resided on the same side or opposite side of the centrosome axis (Fig. S2A’-A”’). ImmunoFISH/WCP was carried out for samples of commercially available neonatal HUVECs (n=2 vials with 10 individuals pooled represented per vial, Table S1). Chromosomes 17 and 19 were selected based on differences in base pair number, and chromosome structure pertaining to the p-arm (non-acrocentric) (Grimwood et al., 2004; Zody et al., 2006). In addition, chromosome 19 is a gene-dense chromosome that has previously been reported to exhibit abnormal homologous chromosome pairing in Caki-1 renal carcinoma cells and renal oncocytoma (Hua and Mikawa, 2018b; Koeman et al., 2008).

Metaphase cells were identified based on the highly condensed chromosomes aligned in the middle of the cell, with centrosomes positioned at the opposite poles (Walczak et al., 2010). Segregation of homologous chromosomes 17 and 19 was observed in a HUVEC at metaphase (Fig. 1C,G). To determine whether segregation of homologous chromosomes 17 and 19 occurred in multiple HUVECs, we created a 3D overlay as previously described (Hua and Mikawa, 2018b) (Fig. S2 for chromosome 19). The centrosome or x-axis, defined by positive γ-tubulin staining, or the center of DNA volume for each cell was used to establish an origin point (0,0,0) for alignment in the 3D overlay analysis (Fig. S2A’). The position of individual chromosomes was determined by identifying the center of volume of the chromosome paint signal (Fig. S2A’). The 3D overlay of chromosomes 17 and 19 at metaphase demonstrated that multiple HUVECs displayed segregation of homologous chromosomes 17 and 19 along the centrosome axis (Fig. 1D,H; Fig. S2). Mapping of individual positions of the homologous chromosomes 17 and 19 in each HUVEC revealed that one homolog consistently positioned itself on each side of the centrosome axis (Fig. 1E,I; segregation in n=17/17 cells for chromosome 17, segregation in n=33/34 cells for chromosome 19; Fig. S3A,A’). In no case did we see homologous chromosomes 17 or 19 exhibit pairing at 0 µm distance or 0° angular orientation relative to its homolog similar to our previous study (Hua and Mikawa, 2018b) (Fig. 1F,J). These data demonstrated that chromosomes 17 and 19 were consistently antipaired in HUVECs at metaphase. Similar to our previous reports at anaphase, these data indicate that the homologous chromosomes 17 and 19 are organized in a non-random, antipaired configuration (Hua and Mikawa, 2018b).

### Homologous chromosome segregation was lost in small chromosomes 17 and 19 in adult HAECs

To investigate whether the antipairing pattern is present in adult aortic endothelial cells (HAECs), we performed ImmunoFISH/WCP for chromosomes 17 and 19 in single donor-derived HAECs (Table S1). Analyzing individual donor samples offer insights into whether this pattern is lost or conserved in adult aortic endothelial cells (ECs) across the general population. If the antipairing pattern is absent in multiple patients, it may suggest that this loss is consistent across the adult aortic EC population. On the other hand, if the antipairing pattern is lost in some patients but still present in others, it could indicate that the presence or absence is specific to individual patients.

HAECs were sourced from individual deceased organ donor patients of various ages, and purchased commercially (Lonza). Some donors had pathologies such as hypertension (n=2/7 patients). For example, one of the 50-year-old females reported hypertension and glioblastoma, while the other 50-year old-female reported fibromyalgia and acute alcohol-related pancreatitis. The 68-year-old male reported hypertension, hyperlipidemia, and diabetes. The other four remaining patients did not report pathologies. In addition, not all patients were analyzed for each chromosome.

To investigate whether this pattern is present across multiple patients, we analyzed HAECs at metaphase from separate patients (Table S1). For chromosome 17 analysis, HAECs from the 36-year-old female, both 50-year-old females, and 68-year-old male were analyzed (Table S1, n=4 patients). For chromosome 19, HAECs from the 36-year-old female, 50-year-old male, one of the 50-year-old females, and 68-year-old male were analyzed (Table S1, n=4 patients). Between chromosomes 17 and 19, HAECs were analyzed from 5 patients total (Table S1). We selected individuals from different age groups and sexes to determine whether the antipairing pattern is preserved among adult aortic ECs.

To determine whether small chromosomes maintain the spatial segregation of homologs in adult HAECs, we analyzed chromosomes 17 and 19 using ImmunoFISH/WCP. We found that both chromosomes 17 and 19 lose the pattern of one homolog per nuclear hemisphere in HAECs derived from five patients (Fig. 1K-R; segregation in=44/83 cells for chromosome 17, segregation in n=17/30 cells for chromosome 19; Fig. S3B,B’; Tables S1, S2). Quantification analysis revealed average distances of 1.68±1.40 µm s.d., 1.99±1.89 µm s.d. and angular orientations of 27.2°±27.5° s.d., 5.43° ± 4.53° s.d. for homologous chromosomes 17 and 19, respectively (Fig. 1N,R; Tables S3, S4).

Notably, chromosome 17 was found to be abnormally paired in 13 out of 83 cells in three patients, the 36-year-old and 50-year-old females, and 68-year-old male (Fig. 1N; Table S1). The 50-year-old female and 68-year-old male also displayed pairing of chromosome 19 in four out of 30 cells (Fig. 1R). Together, these data show that small chromosomes 17 and 19 lose the spatial segregation and exhibit an increase of 15.0% abnormal pairing (17 out of 113 cells) in adult HAECs as compared to 0% of neonatal HUVECs (0 out of 51 cells) (Fig. 1S). A binomial logistic regression was performed to investigate the factors on the probability of pairing of chromosomes 17 and 19 in both HUVECs and HAECs (*X^2^* (1, n=164)=11.6, *P*<0.001).

However, both chromosomes 17 and 19 are small non-acrocentric chromosomes. To further investigate whether other small chromosomes that are acrocentric also lose the spatial segregation, we next mapped the position of small acrocentric chromosomes to determine if the loss of spatial segregation is shared broadly among the human karyotype.

### Homologous segregation was lost in acrocentric chromosomes 13, 15, 21, and 22 in adult HAECs

Acrocentric chromosomes, including 13, 14, 15, 21, 22, and Y, are small chromosomes with the centromere located near one end of the chromosome, and contain the nucleolus organizing region (Colaco and Modi, 2018; Henderson et al., 1972; Lejeune et al., 1960; Rogers, 2021). To test whether small acrocentric chromosomes also lose the one homolog pattern per nuclear hemisphere in adult HAECs, we selected chromosomes 13, 15, 21, and 22 to analyze.

Our previous study showed that the one homolog per nuclear hemisphere motif is a conserved pattern for acrocentric chromosomes in neonatal HUVECs (Hua and Mikawa, 2018b). Three-dimensional overlays of neonatal HUVECs showed segregation of chromosomes 13, 15, 21, and 22 at metaphase (Fig. 2A-I; Movie 1). Pairing of homologous chromosome 21 was observed in two out of 32 cells (Fig. 2I). These data confirm the homologous segregation of acrocentric chromosomes 13, 15, 21, and 22 in HUVECs, with almost no pairing.

**Fig. 2.**
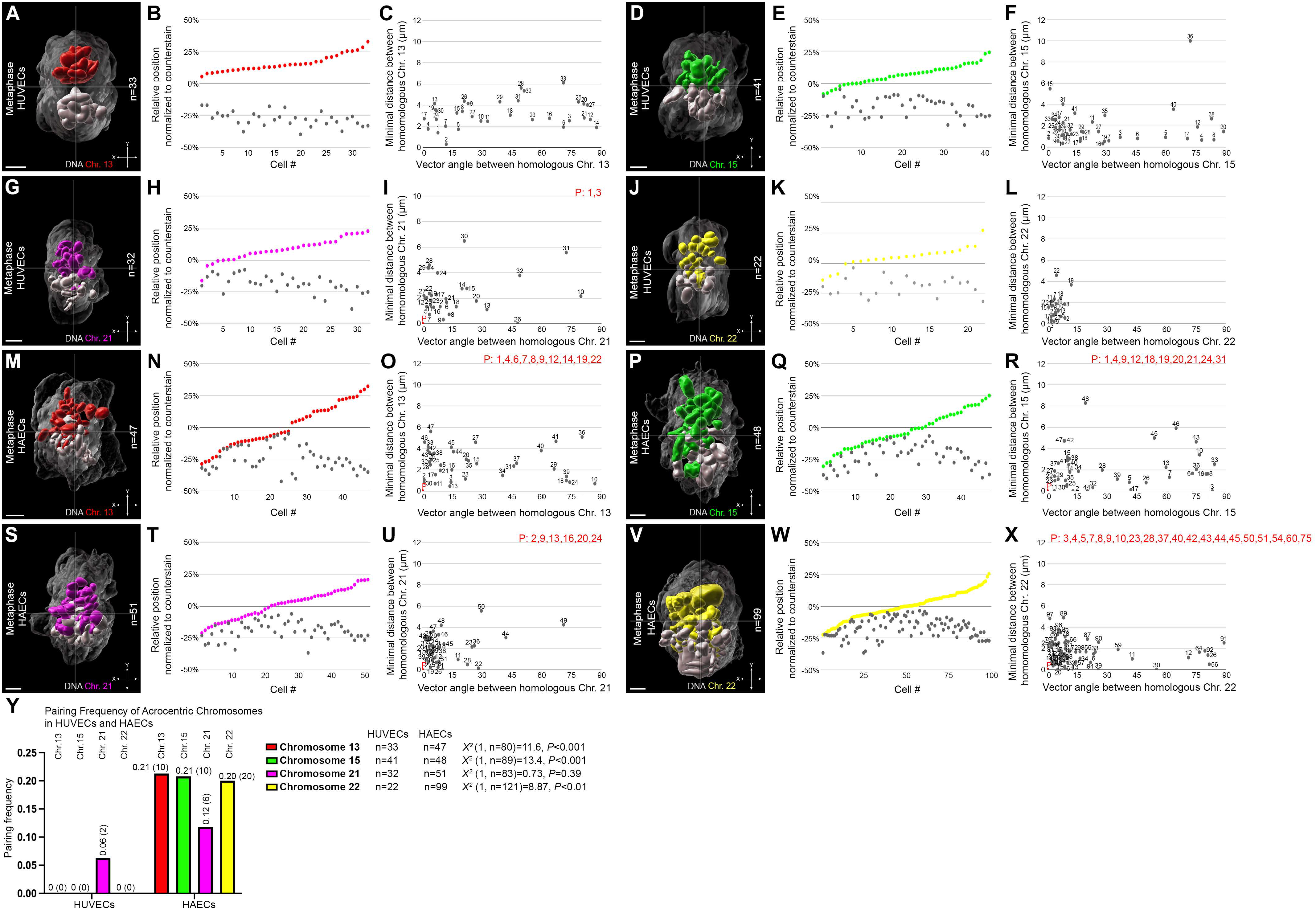
Homologous chromosome segregation was lost in acrocentric chromosomes 13, 15, 21, and 22 in adult HAECs. **(A)** A 3D overlay of chromosomes 13 (red/grey) of multiple neonatal HUVECs at metaphase (segregation in n=33/33 cells). **(B)** Relative positions for each homolog of chromosome 13 (red/grey circles) when mapped to an axial coordinate system. **(C)** The distance and angular orientation of homologous chromosome 13 pairs with an average distance of 3.19±1.27 µm s.d. and angular orientation of 39.7°±29.4° s.d. *Note: Each cell is numbered as in* Fig. 1. **(D-F)** As in (A-C) but of homologous chromosome 15 (green/grey) (segregation in n=33/41 cells). The distance and angular orientation of chromosome 15 homologs with an average distance of 1.88±1.72 µm s.d. and angular orientation of 27.5°±27.7° s.d. **(G-I)** As in (A-C) but of chromosome 21 homologs (magenta/grey) (segregation in n=27/32 cells). The distance and angular orientation of chromosome 21 homologs with an average distance of 2.21±1.60 µm s.d. and angular orientation of 15.7°±20.3° s.d. **(J-L)** As in (A-C) but of chromosome 22 homologs (yellow/grey) (segregation in n=18/22 cells). The distance and angular orientation of chromosome 22 homologs with an average distance of 1.50±1.09 µm s.d. and angular orientation of 4.56°±2.73° s.d. **(M-O)** As in (A-C), but of adult HAECs (n=4 patients, segregation in n=22/47 cells) with an average distance of 2.06±1.66 µm s.d. and angular orientation of 20.8°±26.6° s.d. **(P-R)** As in (D-F), but of adult HAECs (n=3 patients, segregation in n=19/48 cells). The distance and angular orientation of chromosome 15 homologs with an average distance of 1.67±1.85 µm s.d. and angular orientation of 26.6°±29.3° s.d. **(S-U)** As in (G-I), but of adult HAECs (n=4 patients, segregation in n=30/51 cells) with an average distance of 1.81±1.35 µm s.d. and angular orientation of 8.78°±12.5° s.d. **(V-X)** As in (J-L), but of adult HAECs (n=7 patients, segregation in n=48/99 cells). The distance and angular orientation of homologues of chromosome 22 with an average distance of 1.28±1.13 µm s.d. and angular orientation of 12.6°±20.0° s.d. **(Y)** Pairing frequency graph showing the pairing of chromosomes 13 (red) *X^2^* (1, n=80)=11.6, *P*<0.001, 15 (green) *X^2^* (1, n=89)=13.4, *P*<0.001, 21 (magenta) *X^2^* (1, n=83)=0.73, *P*=0.39, 22 (yellow) *X^2^* (1, n=121)=8.87, *P*<0.01 of HUVECs and HAECs. Pairing data were analyzed using binomial logistic regression. Scale bar: 2 µm.

In contrast, 3D overlays of adult HAECs for chromosomes 13, 15, 21, and 22 demonstrated a loss of the one homolog per nuclear hemisphere motif (Fig. 2M-X; Table S2). Abnormal pairing of the acrocentric chromosomes 13, 15, 21, and 22 was observed in 4 out of 4 patients, 2 out of 3 patients, 3 out of 4 patients, and 3 out of 7 patients, respectively (Fig. 2M-X; Table S1). Together, these data show that small acrocentric chromosomes 13, 15, 21, and 22 lose spatial segregation and exhibit a 18.8% increase of abnormal pairing (46 out of 245 cells) in adult HAECs as compared to 1.56% of neonatal HUVECs (2 out of 128 cells) (Fig. 2Y; Table S2; *X^2^* (1, n=373)=30.7, *P*<0.0001). The one homolog per nuclear hemisphere motif along the centrosome axis was lost for the acrocentric chromosomes 13, 15, 21, and 22 in seven adult patients.

These data show that small chromosomes, both non-acrocentric and acrocentric, lose the one homolog per nuclear hemisphere motif and exhibit an increased frequency of homologous pairing. To rule out the possibility that a loss of antipairing and an increase in abnormal pairing events are not limited to small chromosomes only, we next investigated positioning of large chromosomes.

### Fewer adult HAECs showed segregation for the larger chromosomes 1 and 4 as compared to neonatal HUVECs

To test whether large chromosomes also lose the spatial segregation of homologous chromosomes, as observed in non-acrocentric and acrocentric adult HAECs, we mapped the positions of chromosomes 1 and 4 using ImmunoFISH/WCP. Compared to chromosomes 1 and 4 in HUVECs at metaphase, which showed consistent segregation along the centrosome axis (Fig. 3A-F), fewer HAECs displayed segregation of the larger chromosomes 1 and 4 (Fig. 3G-L) (n=5 patients for each chromosome 1 and 4, respectively; Table S2). These findings indicate a gradual loss of fidelity in the spatial segregation of homologous chromosomes in adult aortic ECs.

**Fig. 3.**
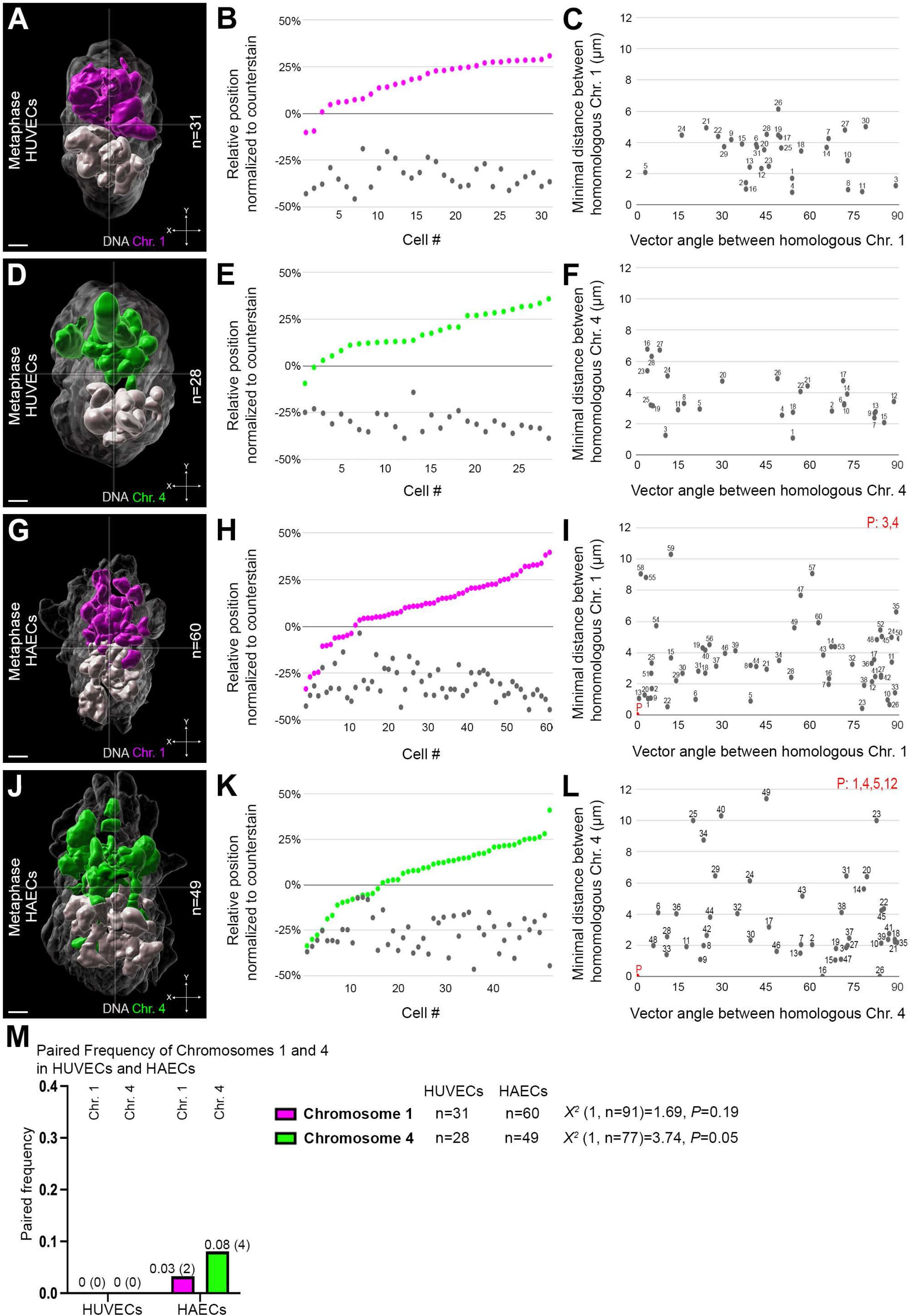
Fewer adult HAECs showed segregation for the larger chromosomes 1 and 4 as compared to neonatal HUVECs. **(A)** A 3D overlay of chromosome 1 (magenta/grey) of multiple neonatal HUVECs at metaphase (segregation in n=29/31 cells). **(B)** Relative positions for each homolog of chromosome 1 (magenta/grey circles) when mapped to an axial coordinate system. **(C)** The distance and angular orientation for the chromosome 1 homologs with an average distance of 3.25±1.45 µm s.d. and angular orientation of 48.2°±19.3° s.d. *Note: Each cell is numbered as in* Fig. 1. **(D-F)** As in A-C, but of chromosome 4 (segregation in n=26/28 cells). Chromosome 4 homologs have an average distance of 3.67±1.48 µm s.d. and angular orientation of 43.9°±30.8° s.d. **(G-I)** As in A-C, but of adult HAECs. The distance and angular orientation chromosome 1 homologs (n=6 patients, segregation in n=48/60 cells) with an average distance of 3.45±2.30 µm s.d. and angular orientation of 45.8°±32.3° s.d. **(J-L)** As in D-F, but of adult HAECs (n=5 patients, segregation in n=34/49 cells). The distance and angular orientation of chromosome 4 homologs with an average distance of 3.38±2.88 µm s.d. and angular orientation of 48.6°±30.1° s.d. **(M)** Pairing frequency graph of chromosome 1 (magenta) *X^2^* (1, n=91)=1.69, *P*=0.19 and chromosome 4 (green) *X^2^* (1, n=77)=3.74, *P*=0.05 for both HUVECs and HAECs. Pairing data were analyzed using binomial logistic regression. Scale bar: 2 µm.

Notably, pairing of homologous chromosomes 1 and 4 was observed in single HAECs of both the 21-year-old and 50-year-old males (Fig. 3I,L). Abnormal pairing of chromosomes 1 and 4 was observed in different cells. Two cells of the 50-year-old female also showed chromosome 4 pairing (Fig. 3L). In contrast, there was no abnormal pairing of chromosomes 1 and 4 in HUVECs (Fig. 3C,F). These data revealed that large chromosomes 1 (3.33%) and 4 (8.16%) in adult HAECs, with a combined 5.50% of cells (6 out of 109), displayed abnormal pairing. This contrasts with neonatal HUVECs, where no abnormal pairing was observed (0 out of 59 cells) (Fig. 3M; *X^2^* (1, n=168)=4.97, *P*=0.026). Next, we investigated whether the sex chromosomes also exhibit a loss of spatial segregation and abnormal pairing in adult HAECs.

### Female adult HAECs showed an increase in abnormal pairing of XX chromosomes as compared to XY chromosomes in male HAECs

To test whether the X chromosomes in females or the XY chromosomes in males lose their segregation pattern in adult HAECs, we performed ImmunoFISH/WCP (Fig. 4G-L). HUVECs have been reported to show spatial segregation of the XX and XY sex chromosomes along the centrosome axis in female and male neonatal HUVECs supporting that the mitotic antipairing organization is sex-type independent (Hua and Mikawa, 2018b). The XX and XY chromosomes in female and male HUVECs showed segregation (Fig. 4A-F) consistent with our previous results (Hua and Mikawa, 2018b). In no case did we observe homologous XX or XY chromosomes exhibit pairing at 0 µm distance or 0° angular orientation relative to its partner (Fig. 4C,F).

**Fig. 4.**
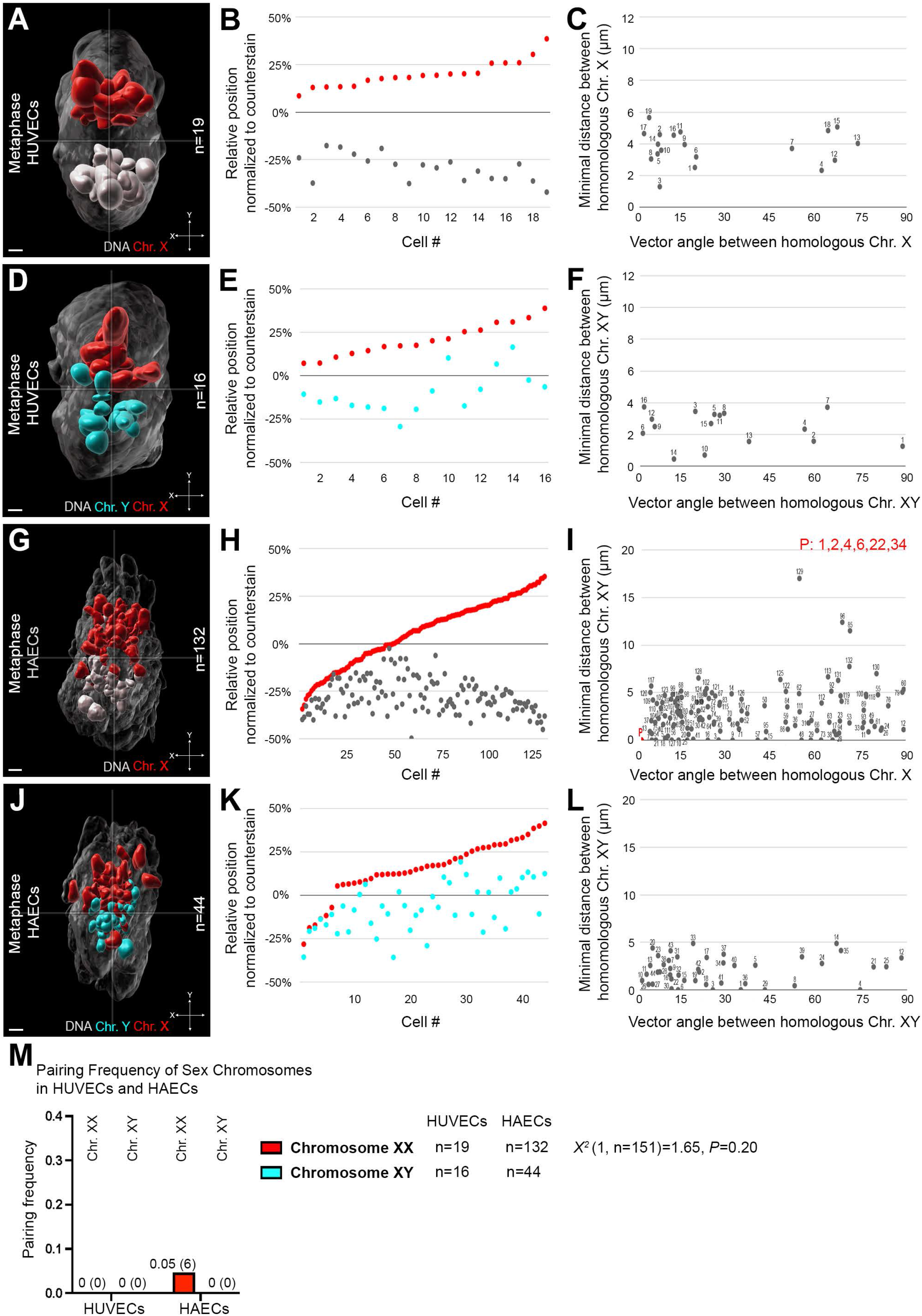
Female adult HAECs showed an increase in abnormal pairing of XX chromosomes as compared to XY chromosomes in male HAECs. **(A)** A 3D overlay of chromosome XX (red/grey) of multiple neonatal HUVECs at metaphase (segregation in n=19/19 cells). **(B)** Relative positions for each homolog of chromosome X (red/grey circles) when mapped to an axial coordinate system. **(C)** The distance and angular orientation of chromosome X homologs with an average distance of 3.79±1.08 µm s.d. and angular orientation of 26.5°±26.7° s.d. *Note: Each cell is numbered as in* Fig. 1. **(D-F)** As in A-C, but of XY (red/cyan) sex chromosomes (segregation in n=13/16 cells) with an average distance of 2.42±1.06 µm s.d. and angular orientation of 29.5°±25.5° s.d. **(G-I)** Same as A-C, but of female-derived HAECs (n=4 patients, segregation in n=82/132 cells). The distance and angular orientation of chromosome X homologs with an average distance of 2.79±2.57 µm s.d. and angular orientation of 35.2°±26.7° s.d. **(J-L)** As in D-F, but of multiple male-derived HAECs (n=3 patients, segregation in n=22/44 cells) with an average distance of 2.04±1.44 µm s.d. and angular orientation of 27.2°±24.6° s.d. **(M)** Pairing frequency graph of chromosome XX (red), *X^2^* (1, n=151)=1.65, *P*=0.20, and chromosome XY (cyan) of HUVECs and HAECs. Pairing data were analyzed using binomial logistic regression. *Note: Statistical analysis could not be performed on chromosomes XY since no pairing was observed for both HUVECs and HAECs.* Scale bar: 2 µm.

However, female adult HAECs showed a loss of X chromosome segregation, and an increase in abnormal pairing as compared to HUVECs (Fig. 4G-I; Table S2; *X^2^* (1, n=151)=1.65, *P*=0.20). Six cells from the 36-year-old, 45-year-old, and 50-year-old female patients showed abnormal pairing of 4.55% (n=6/132 cells). These data suggest that homologous chromosomes X, as well as the large chromosomes 1 and 4, exhibit similar abnormal pairing frequencies in adult HAECs.

Acrocentric autosomes in adult HAECs are shown to be abnormally paired and lose their segregation (Fig. 2M-X). The Y chromosome is also an acrocentric chromosome (Colaco and Modi, 2018; Rogers, 2021). To test whether the XY chromosomes also lose the one sex chromosome per nuclear hemisphere motif in adult HAECs, we performed ImmunoFISH/WCP of three male donor-derived HAECs (Fig. 4J; Table S1). We found that XY chromosomes lose the one sex chromosome per hemisphere motif (Fig. 4J-L, n=44 cells; Table S2). Pairing of XY was not observed in contrast to the abnormal pairing observed in XX in female adult HAECs (Fig. 4I,L,M). These data suggest that there may be a greater affinity for chromosomes with similar sequence homology like the X chromosomes or homologous chromosomes to be abnormally paired as compared to heterologous XY chromosomes. Taken together, X chromosomes in HAECs showed an increased frequency of abnormal pairing as compared to XY chromosomes, with no pairing.

Collectively, small, large, and the XX chromosomes showed a higher frequency of abnormal pairing in adult HAECs compared to neonatal HUVECs (Fig. 5A; *X* ^2^ (1, n=856)=42.5, *P*<0.0001).

**Fig. 5.**
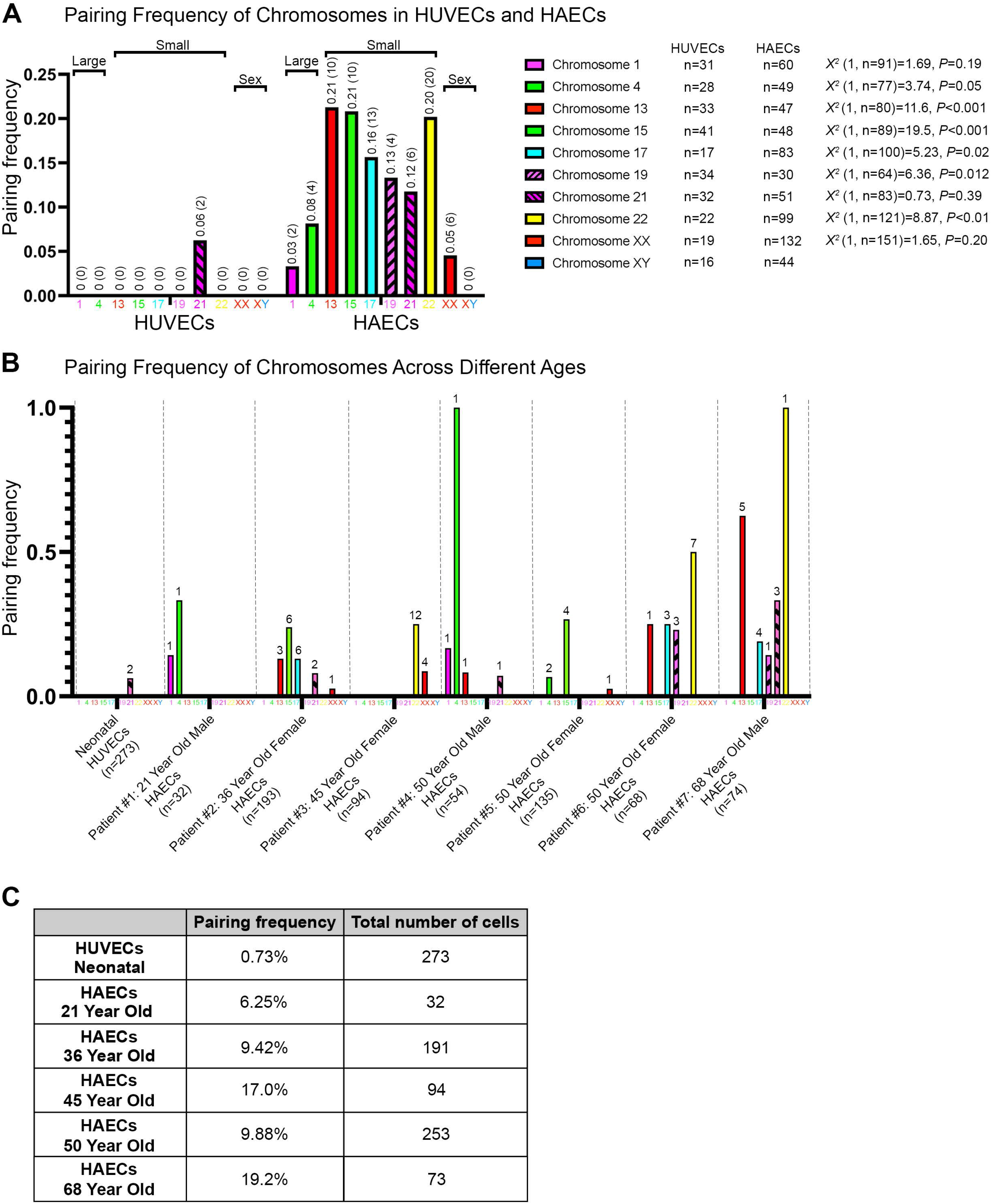
Pairing frequency analysis showed smaller chromosomes had an increase in abnormal pairing as compared to larger chromosomes in adult HAECs. **(A)** Chart representing pairing frequencies of HUVECs and HAECs across chromosomes 1 (magenta), 4 (green), 13 (red), 15 (green), 17 (cyan), 19 (magenta, fill pattern left), 21 (magenta fill pattern right), 22 (yellow), XX (red), and XY (blue). The frequency of homologous chromosome pairing varies among the chromosomes analyzed (*X^2^* (9, n=916)=36.4, *P*<0.0001), for all chromosomes individually of HUVECs and HAECs (for chromosomes individually of HAECs (*X^2^* (9, n=643)=39.2, *P*<0.0001)). For all chromosomes based on sizing of small, large, and sex chromosomes of HUVECs and HAECs (*X^2^* (2, n= 916)=26.1, *P*<0.0001). Pairing data were analyzed using binomial logistic regression. **(B)** Chart of pairing frequency of chromosomes across different age groups. *Note: Numbers above bars indicate the number of pairing events for each chromosome.* **(C)** Table of pairing frequency percentage and sample sizes of analyzed cells across different age groups.

### Pairing frequency analysis showed smaller autosomes had an increase in abnormal pairing as compared to larger chromosomes and sex chromosomes in adult HAECs

It has been proposed that the loss of the antipairing organization can increase the likelihood of somatic homologous chromosome pairing, and mitotic recombination, due to the close proximity of the homologs (Hua and Mikawa, 2018b). The loss of segregation leads to an increased frequency of abnormal pairing observed for chromosomes 1, 4, 13, 15, 17, 19, 21, 22, and XX, with the exception of XY sex chromosomes in adult HAECs (Fig. 5A). While fewer HAECs within the HAEC samples showed a loss of spatial segregation of the large chromosomes 1, 4, and XX in females, this loss was still detectable (Table S2).

Smaller autosomes were observed to have an increase in abnormal pairing as compared to larger autosomes, and sex chromosomes in HAECs (*X* ^2^ (2, n= 643)=31.3, *P*<0.0001). For example, chromosomes 1, 4, and XX which are larger, showed less pairing (4.98%, 12 out of 241 cells), as compared to smaller chromosomes 13, 15, 17, 19, 21, and 22 which exhibited increased abnormal pairing (17.6%, 63 out of 358 cells) (Fig. 5A). In addition, acrocentric chromosomes compared to non-acrocentrics also had an increased frequency of pairing events (*X* ^2^ (1, n=643)=9.18, *P*=0.0024). Our data suggest the underlying mechanisms of homologous segregation and pairing differentially impact chromosomes of different sizes of HAECs (Table S2).

## Discussion

The spatial separation of homologous chromosomes along the centrosome axis, or antipairing, likely plays a fundamental role in maintaining nuclear organization and genome stability. Beyond preventing mitotic recombination, antipairing may serve several critical functions, including preserving nuclear architecture, minimizing inappropriate interchromosomal interactions, ensuring the fidelity of chromosome segregation during mitosis, and supporting precise gene expression programs. Disruption of this organization could therefore have significant consequences for cellular function and genomic integrity.

Previous work showed that neonatal ECs display spatial segregation of homologous chromosomes during metaphase/early anaphase (Cai, Casas, Quintero Plancarte et al., 2025; Hua and Mikawa, 2018b). Here, we find that in adult aortic ECs, the spatial segregation of homologous chromosomes is lost for small chromosomes, and the XY sex chromosomes. Notably, segregation and pairing frequencies vary by chromosome size in HAECs, with smaller chromosomes showing a loss of segregation, and a higher frequency of pairing compared to large chromosomes (Fig. 5A; Table S2). The loss of spatial segregation leads to an increased frequency of pairing of chromosomes 13, 15, 17, 19, 21, 22, with the exception of XY sex chromosomes, in HAECs (Fig. 5A; Table S2). While fewer adult HAECs showed a loss of segregation of chromosomes 1, 4, and XX, there was an increased frequency of pairing observed when compared to HUVECs (Fig. 5A, Table S2).

These observations raise the possibility that the loss of antipairing is not uniform across the genome, but may follow structural or positional rules. This differential homolog pairing based on chromosome number and morphology provides mechanistic hints underlying the antipairing organization. These findings suggest that after birth there may be an increased susceptibility to a loss of nuclear organization, impacting in particular small acrocentric chromosomes (Fig. 5B,C).

Consistent with our observations of homologous segregation, prior studies using fixed HeLa cell and human fibroblast analysis at mitosis show a nonrandom, antiparallel arrangement of homologous chromosomes (Nagele et al., 1995, 1998). Although we did not find an antiparallel configuration of homologous chromosomes in our previous study (Hua and Mikawa, 2018b), these prior publications support that a nonrandom chromosome organization pattern such as haploid set segregation is present.

Live-imaging analysis of hTERT-RPE-1 cells report a random distribution of homologous chromosomes (Stamatov et al., 2025). One interpretation is that immortalized cell lines may not display the same antipairing organization as in primary human cells because antipairing mechanisms may not be present.

In the small chromosomes that were analyzed, we found acrocentric chromosomes 13, 15, and 22 had a higher pairing frequency than small, non-acrocentric chromosomes 17 and 19 (Fig. 5A). One possible explanation for the differential behavior of these acrocentric versus non-acrocentric chromosomes is the presence of interchromosomal contacts among acrocentrics (Henderson et al., 1973; Potapova et al., 2019). These chromosomes, which are involved in nucleolus assembly, also contain interchromosomal DNA connections (Lezhava, 1984; Potapova et al., 2019). It is plausible that these interchromosomal connections are disrupted first, destabilizing acrocentric chromosomes and eventually influencing broader chromosomal organization. However, acrocentric chromosome 21 had a lower pairing frequency of 0.12 as compared to other acrocentric chromosomes in HAECs (Fig. 5A). Chromosome 21 also displayed 2 pairing events in HUVECs, suggesting this acrocentric chromosome may behave differently than others (Fig. 5A).

Mitotic recombination events between heterologous acrocentrics have also been reported (Guarracino et al., 2023; Guissani et al., 1996). Moreover, smaller chromosomes tend to localize more centrally within the metaphase plate, along the centrosome axis, than larger chromosomes, suggesting a size-dependent spatial organization (Hua and Mikawa, 2018b; Hua et al., 2022; Sun et al., 2000).

Research has highlighted a link between chromosome organization and aging (Chandra and Kirschner, 2016; Guarente, 1996; Lezhava, 2001; Liu et al., 2022; Naylor and Deursen, 2016). As ECs age, they exhibit decreased proliferative capacity, leading to senescence, impaired wound healing, and a decline in vascular repair (Bhayadia et al., 2016; Bloom et al., 2023a,b; Guo et al., 2022; Jia et al., 2019; Liu et al., 2019; Rippe et al., 2012; Shabanian et al., 2024; Ting et al., 2021). One common feature is aneuploidy, an abnormal number of chromosomes which can impair normal cell function and is associated with vascular aging (Donato et al., 2015, 2018). For example, human aortic ECs derived from older patients 30 years and older show a higher frequency of aneuploidy (such as tetrasomy) compared to ECs from younger donors (18 months to 30 years old) (Aviv et al., 2001).

Small chromosomes, in particular chromosome 13, have been reported to be gained/lost with age, or passage in ECs and in hematological disorders (Aviv et al., 2001; Hamuro et al., 2016; Kimura et al., 2004; Miyai et al., 2008; Zhang et al., 2000). This pattern of instability may also extend to the sex chromosomes (Guttenbach et al., 1995; Hando et al., 1994; Martin et al., 1980; Miyai et al., 2008; Nowinski et al., 1990; Richard et al., 1994; Stone and Sandberg, 1995). The Y chromosome has been shown to exhibit age-related loss in several tissues (Aviv et al., 2001; Catalán et al., 1998; Gutiérrez-Hurtado et al., 2024; Nath et al., 1995; Pierre and Hoagland, 1972; Sakurai and Sandberg, 1976; Wiktor et al., 2000; Zhang et al., 2007). While the XY sex chromosomes lose the segregation pattern, they do not show increased pairing events like the autosomes (Fig. 4J-L; Fig. 5A; Table S2). This difference likely reflects their unique sequence properties and structural organization. The XY chromosomes share only limited sequence homology, primarily in the pseudoautosomal regions (PAR1/2 regions) (Helena Mangs and Morris, 2007), and undergo independent pairing mechanisms distinct from those of homologous autosomes in both developmental and aging processes (Gutiérrez-Hurtado et al., 2024; Jacobs et al., 2012; Nath et al., 1995; Weng et al., 2016). Consistent with this, the Y chromosome also undergoes frequent loss in aging tissues, particularly in the hematopoietic and blood cell lineages (Aviv et al., 2001; Gutiérrez-Hurtado et al., 2024; Jacobs et al., 1963; Miyai et al., 2008; Nath et al., 1995; Pierre and Hoagland, 1972; Sakurai and Sandberg, 1976; Stone and Sandberg, 1995; Weng et al., 2016; Wiktor et al., 2000; Zhang et al., 2007). It may be uniquely vulnerable to genomic instability due to its structure, limited recombination, and small size (Colaco and Modi, 2018; Rogers, 2021).

Due to the challenges in culturing human adult aortic ECs, we were only able to analyze samples from seven adult organ donor patients: One male each aged 21, 50, and 68, two females aged 36 and 45, and two females aged 50 (Table S1). As a result, we cannot exclude the possibility that these individual cases may not reflect the broader population for chromosome organization (Lau et al., 2020).

We observed abnormal pairing of homologous chromosomes for all seven patients, rather than only a few. This suggests the loss of antipairing is a common feature in adult aortic ECs (Fig. 5C). Whether the loss of antipairing in adult HAECs increases with age, and is associated with sex still remains elusive. Our study analyzed a subset of individuals across different age groups, which limited statistical analysis of age, and age with sex determinants (Table S1). Nonetheless, graphical data suggests a preliminary age-based loss of spatial segregation of homologous chromosomes, with a trend in the increase of pairing frequency during adulthood (Fig. 5B).

There are at least two potential causes for the higher pairing frequency or loss of antipairing: one could be an age-dependent change, while the other may involve an aorta-specific mechanism that enhances the loss of antipairing. These hypotheses, age-related versus tissue-specific, can be independently evaluated. To investigate the potential role of age, future experiments can expand our analysis to include additional HAEC lines from younger donors, particularly 21- and 36-year olds, as our current dataset includes only one individual from each of these age groups. By comparing these additional younger to older individuals, such as the 50-year-old donors in our current study, we can assess whether abnormal pairing frequencies correlate with age and gain statistical power to evaluate this trend. To test for a tissue-specific mechanism, future experiments can examine endothelial cells from a different vascular region, such as human iliac artery endothelial cells (HIAECs), across various age groups. Identifying similar pairing abnormalities in HIAECs would suggest a broader, systemic feature, while their absence would point toward aorta-specific regulation. As this study focused exclusively on adult aortic endothelial populations, it remains important to evaluate additional adult primary human EC types to determine whether antipairing loss is cell-type specific.

Importantly, the spatial arrangement of chromosomes within the cell can become disordered in adult aortic ECs, potentially affecting gene expression and overall cellular function. However, adult ECs are less proliferative (Lau et al., 2020) than neonatal HUVECs, so genomic instability may mitigate the consequences, as fewer cellular divisions occur. Our findings offer a better understanding of the mitotic antipairing pattern observed in human neonatal ECs, which may be lost with age. This loss of higher-order chromosome organization highlights its critical role in maintaining genomic stability in human cells.

## Materials and Methods

### Cell culture

Primary human umbilical vein endothelial cells (HUVECs, ATCC (Manassas, VA, USA): Cat. No. PSC-100-013, Lot # 70032758 and 81006213) were cultured according to established protocols (Cai, Casas, Quintero Plancarte et al., 2025; Hua and Mikawa, 2018b). Each lot consisted of pooled neonatal HUVECs from ten donors, representing a total of 20 individual donor samples across both lots.

Human aortic endothelial cells (HAECs), (Lonza (Basel, CHE): Cat. No. CC-2535, Lot #: 0000430739, 22TL13791, 000225231, 0000351486, 0000337673, 21TL076036, 22TL073027) were derived from individual human donors (21 year old male, 36 year old female, 45 year old female, 50 year old male, two 50 year old females, and 68 year old male) and were fixed at low passage (<7). MCDB-131 (Cat No. 10372-019), 1% Glutamax (Cat No. 35050-061), 1% Pen-strep (Cat No. 15140-122), and 2% Large vessel endothelial supplement (Cat No. A14608-01) were used for cell growth. Cells were grown on a 10 cm dish until reaching a confluency of 70-80% and then transferred onto a 15 cm plate containing 2 well, 8 mm well diameter PTFE glass slides (Electron Microscopy Sciences (Hatfield, PA, USA), Cat. No. 63416-08), which were flamed/UV sterilized.

### Immunofluorescence and Chromosome painting

HUVECs and HAECs were grown on PTFE glass slides to 80-90% confluence. HUVECs and HAECs were not synchronized prior to metaphase analysis to minimize potential confounding effects associated with cell synchronization procedures (Rodman et al., 1980; Rohlf et al., 1980). Cells were fixed with 4% paraformaldehyde to preserve the three-dimensional nuclear and chromosomal architecture as paraformaldehyde fixation is widely recognized for maintaining nuclear integrity (Hepperger et al., 2007; Hobro and Smith, 2017; Hua and Mikawa 2018a,b, 2018; Kozubek et al., 2000). Following fixation, the slides were stored in 70% ethanol at −20°C for a minimum of 24 hours. Imaging was performed immediately after completion of the experimental procedures, following curing of the samples in ProLong Antifade mounting medium, according to manufacturer’s instructions.

### Immunofluorescence

Slides were washed three times in 4°C 1x Phosphate Buffered Saline (PBS) on ice before undergoing heat-induced antigen retrieval for 10 minutes in a sodium citrate buffer solution (10 mM sodium citrate, 0.05% Tween-20, pH 6.0) in a steamer. Slides were then washed in a permeabilization buffer (0.25% Triton X-100 in 1x PBS) for 10 minutes followed by three 1x PBS washes at room temperature (RT). A blocking buffer (10% goat serum, 0.1% Tween-20 in 1x PBS) was then applied for 30 minutes at RT. Slides were incubated with a primary antibody to γ-tubulin [1:1000, abcam: Rabbit Anti-γ-tubulin antibody (Abcam (Cambridge, GBR) Cat# ab11317, RRID:AB_297921)] diluted in 10% goat serum in 1x PBS with 0.1% Tween-20. Parafilm was used to cover slides. Slides were then incubated at 4°C in a humidified chamber overnight. The following day, the slides were washed three times with 1x PBS. Incubation of a secondary antibody [1:500, abcam: Goat Anti-Rabbit IgG H&L (Alexa Fluor® 647) (Abcam Cat# ab150079, RRID:AB_2722623)] (in 1% BSA in 1x PBS with 0.1% Tween-20) was applied to slides, covered in Parafilm, and incubated in the dark for 1 hour at RT. Slides were then washed twice with 1x PBS, and processed with DNA counterstain DAPI (Invitrogen: D1306) for immunofluorescence. The slides were washed twice in 1x PBS. After, Antifade (Invitrogen (Waltham, MA, USA): Cat. No. P36930) was applied with a 24X60 mm coverslip (Corning (Corning, NY, USA): Cat. No. 2980-246, #1.5) and sealed with nail polish. The slides were stored at 4°C until using the confocal microscope.

### Immunofluorescence and Chromosome painting

If continuing on for chromosome painting, before processing with the DNA counterstain steps listed above, the slides were incubated with an EGS crosslinker solution (25% DMSO, 0.375% Tween-20, 25mM EGS (Thermo Scientific: Cat. No. 21565) in 1x PBS) for 10 minutes in the dark at RT. Slides were washed twice with 1x PBS. Whole Chromosome Paints for chromosomes 1, 4, 13, 15, 17, 19, 21, 22, X, and Y in Aqua, Texas red, or FITC (Applied Spectral Imaging, ASI (Carlsbad, CA, USA)) were used to visualize individual chromosomes in metaphase cells.

Chromosome paint probes (ASI) are generated using flow-sorted chromosomes. DOP-PCR is used for DNA amplification with a semi-degenerate primer that anneals at random positions across the genome, with a bias towards primer binding site regions with more compatible 6-mer sequences. Thousands of DNA fragments are generated for each chromosome probe, ranging in size from approximately 200 bp to ∼2,000 bp, with a peak distribution around 400–800 bp. The coverage of these fragments is broad but not uniform, with an estimation of ∼70–85% coverage of the whole chromosome and similar for all chromosomes. Repetitive sequences such as centromeres and telomeres are underrepresented as they are blocked by Cot-1 DNA. The rDNA and X/Y Homologous Sequences can also display cross-hybridization artifacts and non-specific signals on the acrocentric and sex chromosomes.

The probes were preheated on an 80°C heat block for 10 minutes. Then, the probes were incubated at 37°C for 1 hour. The slides were washed with a buffer containing saline-sodium citrate (2x SSC) (Cat. No. 15557-044). After, the slides were submerged in 0.1 N HCl for 5 minutes at RT. Then, the slides were washed with 1x PBS, three times for 5 minutes, followed by a series of chilled ethanol washes on ice. The slides were placed in a denaturation solution (20X SSC, formamide, deionized water) for 7 minutes at 80°C followed by a second series of cold ethanol washes on ice. The slides were incubated with probes on a slide warmer. An 8 mm (World Precision Instruments (Sarasota, FL, USA): Cat. No. 052041) and 12 mm coverslip (Electron Microscopy Sciences: Cat. No. 72290-04) were stacked in subsequent order on the wells, and sealed with rubber cement. The slides were incubated in a moisturizing container containing formamide and deionized water, overnight at 37°C. The following day, the slides were washed in a post-hybridization buffer (1% Tween-20, 10% 20x SSC in deionized water) at 42°C for 10 minutes, three times. Slides were then incubated with a DNA counterstain (DAPI, Hoechst, SYTOX, or TO-PRO-3, Invitrogen: Cat. No. D1306, H1399, S11380, T3605). The slides were washed twice in 1x PBS. After, Antifade was applied with a 12 mm coverslip and sealed with nail polish. The slides were stored at 4°C.

### Confocal microscopy

Fixed cells were imaged using a laser scanning confocal microscope (Leica (Wetzlar, DEU): TCS SPE, DM2500) using a 63x/1.3NA oil immersion objective with a digital zoom of 1.5x to visualize individual cells. The z-stacks (z-step size: 0.35 µm) were acquired sequentially in a four-channel mode; z-stacks were captured using a frame size of 1,024×1,024 pixels and processed with Leica Application Suite X software (Version: 3.5.2.18963).

### 3D reconstruction and overlay of homologous chromosomes

Confocal optical sections were reconstructed in a 3D software, Imaris (Bitplane: 9.7.2). Three-dimensional overlay generation of homologous chromosomes in multiple fixed metaphase HUVECs and HAECs was performed as previously described (Hua and Mikawa, 2018b). We detected a small population of HAECs to be polyploid, consistent with prior reports (Aviv et al., 2001; Borradaile and Pickering, 2010). For example, we found 36 of approximately 207,000 HAECs to be polyploid, and these cells were not included in the analysis.

To distinguish homolog pairing from chromosome loss (monosomy), we quantified chromosome paint signal volume. The volume of a paired chromosome paint signal was approximately equivalent to the combined volume of two individual homologs. For example, the average volume of an individual, antipaired, chromosome 19 homolog paint signal was measured to be 2.15±1.02 µm^3^ s.d. in 10 HUVECs and 2.32±1.08 µm^3^ s.d. in 10 HAECs. However, the average volume of the single chromosome paint signals, in the four cases of chromosome 19 pairing in HAECs, was 5.12±3.38 µm^3^ s.d. These average volumes showed that the paired chromosome 19 paint signals in HAECs were approximately twice the volume of the individual, antipaired, homologous chromosome 19 paint signals in HUVECs and HAECs respectively.

Multiple pairs of homologous chromosomes were visualized by creating the cellular axis that was previously defined by Hua and Mikawa (2018b). The x, y, and z axes were set an origin of 0,0,0 to define each nuclear hemisphere across the x-axis (centrosome axis), and to analyze how each pair of homologous chromosomes spatially position themselves in relation to the centrosome axis. The center of volume of the centrosomes by γ-tubulin immunofluorescence staining defined the x-axis and was used for alignment of metaphase cells.

### Defining the relative position of homologous chromosomes

The centers of volume for the 3D reconstructed surfaces of homolog chromosome paint signals and DNA counterstains were obtained via the statistics tool in Imaris. 3D reconstructed homologs were color-coded as either colored (cyan, magenta, red, yellow, or green) or grey depending on their distance from the x-axis (y=0). The homolog that had the y-value closest to y=0 would be designated as the colored homolog and the other respective homolog would be grey. Furthermore, the cells were oriented so that colored homologs occupied the more positive positions than the grey homologs. The position of each homolog was analyzed using the “Position Y” statistic, which provided the y-coordinate of the center of volume of the 3D surface.

The position of each pair of homologous chromosomes was normalized in every mitotic cell with regards to the DNA counterstain similar to Hua and Mikawa (2018b). The relative positions of each homologous chromosome was calculated by normalizing the position of the y-coordinate of the homolog to the total length of the DNA metaphase plate along the y-axis (BoundingBoxAA Length Y) according to the following equation:

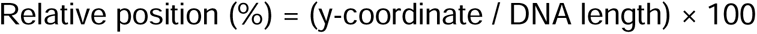

The normalization scales chromosome positions to a range of approximately −50% to +50%, corresponding to the lower and upper halves of the metaphase plate, respectively. For example, a chromosome 19 homolog in HUVECs had a y-coordinate of 2.71 µm and a DNA metaphase plate length of 9.12 µm. The relative position was therefore calculated as (2.71/9.12) × 100 = 29.7%.

Immunofluorescence staining for γ-tubulin, along with a DNA counterstain, revealed that the centrosome axis, and the volume-derived axis positioned perpendicular to the longest axis of the metaphase DNA plate, with an average distance of 0.5 µM apart (Fig. S1). Consequently, individual chromosome positions within ± 0.5 µM of the DNA center of volume, referred to as the “boundary region”, were excluded from the analysis. For instance, out of 66 total cells, 6 cells exhibited homologous chromosome 1 in this boundary region, rendering the data uninterpretable (n=6/66 cells for chromosome 1).

### 3D distance and angular orientation measurements of homologous chromosomes

Homologous chromosomes were identified as paired if the distance between them was 0 µm (≤0.36 µm, the spatial xy resolution limit of the confocal microscope) and an angular orientation (vector angle) of 0°.

Minimal distances between homologs were determined by the shortest distances between the edges of the 3D surfaces of homologs.

The 3D vector coordinates determined the angular orientation of the homologs. To determine a 3D vector coordinate, a 3D surface of a homolog was selected. The longest ellipsoid axis length (Ellipsoid Axis C) was identified in the “specific values” tab. The x, y, and z coordinates were then acquired to define the 3D vector. This process was repeated for the other homolog. The vector angle was calculated by taking the inverse cosine of the normalized dot product.

At metaphase, it was difficult to determine the centromere position in relation to the chromosome arm due to observing a fixed image when centromeres oscillate along the centrosome axis and chromosome arms are more dynamic. Therefore, homolog orientation was quantified using a single quadrant (90° range) instead of a 180° range that was previously used for 3D vector angle analysis of anaphase cells in which the vectors had a defined direction (Hua and Mikawa, 2018b).

### Statistical analysis

The loss of spatial segregation was used to describe the position of individual homologs across the centrosome axis. The loss of antipairing was used to describe the loss of haploid set organization. Throughout the manuscript, we used these terms to describe the loss of statistically significant non-random spatial segregation, rather than the complete elimination of homolog separation. Our definition was based on statistical testing against the null hypothesis that homologous chromosomes were positioned randomly with respect to one another, similar to nonhomologous chromosome pairs (Hua and Mikawa, 2018).

Pairing was defined as occurring in cells where homologs had a distance and angular orientation difference of 0. Contingency table analysis or binomial logistic regression was used to assess the effect of explanatory factors on the probability of pairing. Statistical analysis was completed using the statistical analysis software, JMP Pro 18 (SAS Institute, Raleigh, NC)).

### Data Visualization

All figures were generated using Adobe Photoshop (Version 26.3.0). Bar graphs depicting pairing frequencies of chromosomes between HUVECs and HAECs were generated with Graphpad Prism 11 (Version 11.0.2 (92)).

Rotational video of both single whole chromosome painting and 3D overlays for HUVECs and HAECs were generated using the video editing software, Adobe Premiere Pro (Version 25.0.0, Build 61). To show side-by-side comparison of chromosome positioning of both HUVECs and HAECs, each cell was repositioned and resized using the position tool for both X and Y, and the scale function within the properties tab. The text was implemented in the video using the type tool within the tools tab and the corresponding text colors were changed within the properties tab. Video was exported and used the preset: Match Source - Adaptive High Bitrate, Format of H.264 and recorded at 24 fps.

## Author Declarations

### Funding

This work was supported in part by National Science Foundation (NSF RUI Award #2027746 to L.L.H.), National Institutes of Health (NIH) (NIGMS Award #R16GM153517 to L.L.H.), California State University Program for Education and Research in Biotechnology (CSUPERB New Investigator Grant, CSUBIOTECH Research Development Grant to L.L.H., and Faculty Graduate Student Collaborative Grant to O.A./L.L.H.), Koret Foundation/ORSP grant (to G.Q.P., J.M., and L.L.H.), and SSU start up funds (to L.L.H.).

### Conflicts of interest/Competing interests

The authors declare no conflicts of interest nor competing interests.

### Ethics approval/declarations

The following study did not involve human participants. Cell lines in the study were purchased through commercial vendors (American Type Culture Collection (ATCC), and Lonza).

### Consent to participate

Not applicable

### Consent to publication

Not applicable

### Availability of data and material/Data availability

All data supporting the findings of this study are available within the paper and its Supplementary Information.

### Code availability

Not applicable

## Author’s contributions

J.M., O.A., and G.Q.P., performed experiments; J.M., O.A., G.Q. P., and L.L.H. conducted analysis; J.M., G.Q.P., and L.L.H. wrote the first draft, and O.A. completed revisions of the manuscript. All authors read and approved the final manuscript.

## Acknowledgements

J.M. would like to express her heartfelt gratitude to her parents, Adolfo Morales and Silvia Cienfuegos, for their unwavering love and support throughout this journey. To her brothers, Adolfo and Alexis, for their encouragement and understanding. J.M. is also deeply grateful to her family friend, Alfonso Ventura, for his invaluable support. His presence and guidance have been instrumental in the completion of her thesis. O.A. would like to express his gratitude to his parents Naheem Akhtar and Kairi Kaarus for their love and support, and his siblings Hannah, Anwar, and Salma for being a steadfast source of optimism and inspiration. G.Q.P. would like to express his gratitude to his brother Christian D. Quintero and his parents Marisela Plancarte Maravilla and Gabriel Quintero Ortega for their love and support. Thank you to Drs. Takashi Mikawa and Joseph Lin for review and discussion of the manuscript, Dr. Daniel Crocker for support and discussion on statistical analysis. Special thanks to the members of the Hua lab, including Emerson Herrmann, for their valuable feedback, guidance, and ongoing support. L.L.H. dedicates this work to the memory of her daughter, Josephine Jereb. Part of this work is incorporated into two Master’s theses (J.M., O.A.).

## Notes

### Competing Interest Statement

The authors have declared no competing interest.

### Summary of Updates

These revisions include the addition of new data (Fig. 4, Supplemental Fig. 3, and Supplemental Tables 2-4), clarification of the analyses, and revisions throughout the text to improve accuracy, and transparency.

## References

Atkin, N. B. and Jackson, Z. (1996). Evidence for somatic pairing of chromosome 7 and 10 homologs in a follicular lymphoma. Cancer Genet. Cytogenet. 89, 129–31. doi:10.1016/0165-4608(95)00360-6

Aviv, H., Khan, M. Y., Skurnick, J., Okuda, K., Kimura, M., Gardner, J., Priolo, L. and Aviv, A. (2001). Age dependent aneuploidy and telomere length of the human vascular endothelium. Atherosclerosis 159, 281–287. 10.1016/s0021-9150(01)00506-8

Bhayadia, R., Schmidt, B. M., Melk, A. and Hömme, M. (2016). Senescence-Induced Oxidative Stress Causes Endothelial Dysfunction. J. Gerontol. A Biol. Sci. Med. Sci. 71, 161–169. 10.1093/gerona/glv008

Bloom, S. I., Islam, M. T., Lesniewski, L. A. and Donato, A. J. (2023). Mechanisms and consequences of endothelial cell senescence. Nat. Rev. Cardiol. 20, 38–51. 10.1038/s41569-022-00739-0

Bloom, S. I., Liu, Y., Tucker, J. R., Islam, M. T., Machin, D. R., Abdeahad, H., Thomas, T. G., Bramwell, R. C., Lesniewski, L. A. and Donato, A. J. (2023). Endothelial cell telomere dysfunction induces senescence and results in vascular and metabolic impairments. Aging cell 22, e13875. 10.1111/acel.13875

Borradaile, N. M. and Pickering, J. G. (2010). Polyploidy impairs human aortic endothelial cell function and is prevented by nicotinamide phosphoribosyltransferase. Am. J. Physiol. Cell Physiol. 298, C66–C74. 10.1152/ajpcell.00357.2009

Burgess, R. C., Misteli, T. and Oberdoerffer, P. (2012). DNA damage, chromatin, and transcription: the trinity of aging. Curr. Opin. Cell Biol. 24, 724–730. 10.1016/j.ceb.2012.07.005

Cai, P., Casas, C. J., Quintero Plancarte, G., Mikawa, T. and Hua, L. L. (2025). Ipsilateral restriction of chromosome movement along a centrosome, and apical-basal axis during the cell cycle. Chromosome Res. 33, 1–21. 10.1007/s10577-024-09760-0

Callaini, G. and Riparbelli, M. G. (1996). Fertilization in *Drosophila melanogaster*: centrosome inheritance and organization of the first mitotic spindle. Dev. Biol. 176, 199–208. 10.1006/dbio.1996.0127

Catalán, J., Autio, K., Kuosma, E. and Norppa, H. (1998). Age-dependent inclusion of sex chromosomes in lymphocyte micronuclei of man. Am. J. Hum. Genet. 63, 1464–1472. 10.1086/302092

Chandra, T. and Kirschner, K. (2016). Chromosome organisation during ageing and senescence. Curr. Opin. Cell Biol. 40, 161–167. 10.1016/j.ceb.2016.03.020

Colaco, S. and Modi, D. (2018). Genetics of the human Y chromosome and its association with male infertility. Reprod. Biol. Endocrinol. 16, 14. 10.1186/s12958-018-0330-5

Cremer, T. and Cremer, C. (2001). Chromosome territories, nuclear architecture and gene regulation in mammalian cells. Nat. Rev. Genet. 2, 292–301. 10.1038/35066075

Cremer, T., Cremer, M., Hübner, B., Silahtaroglu, A., Hendzel, M., Lanctôt, C., Strickfaden, H. and Cremer, C. (2020). The interchromatin compartment participates in the structural and functional organization of the cell nucleus. BioEssays 42, e1900132. 10.1002/bies.201900132

Donato, A. J., Machin, D. R. and Lesniewski, L. A. (2018). Mechanisms of dysfunction in the aging vasculature and role in age-related disease. Circ. Res. 123, 825–848. 10.1161/CIRCRESAHA.118.312563

Donato, A. J., Morgan, R. G., Walker, A. E. and Lesniewski, L. A. (2015). Cellular and molecular biology of aging endothelial cells. J. Mol. Cell. Cardiol. 89, 122–135. 10.1016/j.yjmcc.2015.01.021

Emerson, F. J. and Lee, S. S. (2023). Chromatin: the old and young of it. Front. Mol. Biosci. 10, 1270285. 10.3389/fmolb.2023.1270285

Faruqi, S. A., Miller, R. C. and Noumoff, J. S. (1994). Somatic pairing: an alternative for the development of cancer and other hereditary diseases. Cytologia 59, 439–444. 10.1508/cytologia.59.439

Ghosh, R. P. and Meyer, B. J. (2021). Spatial organization of chromatin: emergence of chromatin structure during development. Annu. Rev. Cell Dev. Biol. 37, 199–232. 10.1146/annurev-cellbio-032321-035734

Gondos, B. and Bhiraleus, P. (1970). Pronuclear relationship and association of maternal and paternal chromosomes in flushed rabbit ova. Z. Zellforsch. Mikrosk. Anat. 111, 149–159. 10.1007/BF00339782

Grimwood, J., Gordon, L. A., Olsen, A., Terry, A., Schmutz, J., Lamerdin, J., Hellsten, U., Goodstein, D., Couronne, O., Tran-Gyamfi, M. et al. (2004). The DNA sequence and biology of human chromosome 19. Nature 428, 529–535. 10.1038/nature02399

Guarente, L. (1996). Do changes in chromosomes cause aging? Cell 86, 9–12. 10.1016/s0092-8674(00)80072-0

Guarracino, A., Buonaiuto, S., de Lima, L. G., Potapova, T., Rhie, A., Koren, S., Rubinstein, B., Fischer, C., Human Pangenome Reference Consortium, Gerton, J. L. et al. (2023). Recombination between heterologous human acrocentric chromosomes. Nature 617, 335–343. 10.1038/s41586-023-05976-y

Guissani, U., Facchinetti, B., Cassina, G. and Zuffardi, O. (1996). Mitotic recombination among acrocentric chromosomes’ short arms. Ann. Hum. Genet. 60, 91–97. 10.1111/j.1469-1809.1996.tb01180.x

Guo, J., Huang, X., Dou, L., Yan, M., Shen, T., Tang, W. and Li, J. (2022). Aging and aging-related diseases: from molecular mechanisms to interventions and treatments. Signal Transduct. Target. Ther. 7, 391. 10.1038/s41392-022-01251-0

Gutiérrez-Hurtado, I. A., Sánchez-Méndez, A. D., Becerra-Loaiza, D. S., Rangel-Villalobos, H., Torres-Carrillo, N., Gallegos-Arreola, M. P. and Aguilar-Velázquez, J. A. (2024). Loss of the Y chromosome: a review of molecular mechanisms, age inference, and implications for men’s health. Int. J. Mol. Sci. 25, 4230. 10.3390/ijms25084230

Guttenbach, M., Koschorz, B., Bernthaler, U., Grimm, T. and Schmid, M. (1995). Sex chromosome loss and aging: in situ hybridization studies on human interphase nuclei. Am. J. Hum. Genet. 57, 1143–1150.

Hamuro, J., Ueno, M., Toda, M., Sotozono, C., Montoya, M. and Kinoshita, S. (2016). Cultured human corneal endothelial cell aneuploidy dependence on the presence of heterogeneous subpopulations with distinct differentiation phenotypes. Invest. Ophthalmol. Vis. Sci. 57, 4385–4392. 10.1167/iovs.16-19771

Hando, J. C., Nath, J. and Tucker, J. D. (1994). Sex chromosomes, micronuclei and aging in women. Chromosoma 103, 186–192. 10.1007/BF00368011

Henderson, A. S., Warburton, D. and Atwood, K. C. (1972). Location of ribosomal DNA in the human chromosome complement. Proc. Natl. Acad. Sci. USA 69, 3394–3398. 10.1073/pnas.69.11.3394

Henderson, A. S., Warburton, D. and Atwood, K. C. (1973). Ribosomal DNA connectives between human acrocentric chromosomes. Nature 245, 95–97. 10.1038/245095b0

Heard, E. and Bickmore, W. (2007). The ins and outs of gene regulation and chromosome territory organisation. Curr. Opin. Cell Biol. 19, 311–316. 10.1016/j.ceb.2007.04.016

Helena Mangs, A. and Morris, B. J. (2007). The human pseudoautosomal region (PAR): origin, function and future. Curr. Genomics 8, 129–136. 10.2174/138920207780368141

Hepperger, C., Otten, S., von Hase, J. and Dietzel, S. (2007). Preservation of large-scale chromatin structure in FISH experiments. Chromosoma 116, 117–133. 10.1007/s00412-006-0084-2

Hobro, A. J. and Smith, N. I. (2017). An evaluation of fixation methods: spatial and compositional cellular changes observed by Raman imaging. Vib. Spectrosc. 91, 31–45.

Hua, L. L. and Mikawa, T. (2018). Chromosome Painting of Mouse Chromosomes. Methods Mol. Biol. 1752, 133–143. 10.1007/978-1-4939-7714-7_13

Hua, L. L. and Mikawa, T. (2018). Mitotic antipairing of homologous and sex chromosomes via spatial restriction of two haploid sets. Proc. Natl. Acad. Sci. USA 115, E12235–E12244. 10.1073/pnas.1809583115

Hua, L. L., Casas, C. J. and Mikawa, T. (2022). Mitotic antipairing of homologous chromosomes. Results Probl. Cell Differ. 70, 191–220. 10.1007/978-3-031-06573-6_6

Jacobs, K. B., Yeager, M., Zhou, W., Wacholder, S., Wang, Z., Rodriguez-Santiago, B., Hutchinson, A., Deng, X., Liu, C., Horner, M. J. et al. (2012). Detectable clonal mosaicism and its relationship to aging and cancer. Nat. Genet. 44, 651–658. 10.1038/ng.2270

Jacobs, P. A., Brunton, M., Court Brown, W. M., Doll, R. and Goldstein, H. (1963). Change of human chromosome count distribution with age: evidence for a sex difference. Nature 197, 1080–1081. 10.1038/1971080a0

Jia, G., Aroor, A. R., Jia, C. and Sowers, J. R. (2019). Endothelial cell senescence in aging-related vascular dysfunction. Biochim. Biophys. Acta Mol. Basis Dis. 1865, 1802–1809. 10.1016/j.bbadis.2018.08.008

Joyce, E. F., Erceg, J. and Wu, C. T. (2016). Pairing and anti-pairing: a balancing act in the diploid genome. Curr. Opin. Genet. Dev. 37, 119–128. 10.1016/j.gde.2016.03.002

Joyce, E. F., Williams, B. R., Xie, T. and Wu, C. T. (2012). Identification of genes that promote or antagonize somatic homolog pairing using a high-throughput FISH-based screen. PLoS Genet. 8, e1002667. 10.1371/journal.pgen.1002667

Keneklis, T. P. and Odartchenko, N. (1974). Autoradiographic visualisation of paternal chromosomes in mouse eggs. Nature 247, 215–216. 10.1038/247215a0

Kimura, M., Cao, X., Patel, S. and Aviv, A. (2004). Survival advantage of cultured human vascular endothelial cells that lost chromosome 13. Chromosoma 112, 317–322. 10.1007/s00412-004-0276-6

Koeman, J. M., Russell, R. C., Tan, M. H., Petillo, D., Westphal, M., Koelzer, K., Metcalf, J. L., Zhang, Z., Matsuda, D., Dykema, K. J. et al. (2008). Somatic pairing of chromosome 19 in renal oncocytoma is associated with deregulated EGLN2-mediated [corrected] oxygen-sensing response. PLoS Genet. 4, e1000176. 10.1371/journal.pgen.1000176

Kozubek, S., Lukásová, E., Amrichová, J., Kozubek, M., Lisková, A. and Slotová, J. (2000). Influence of cell fixation on chromatin topography. Anal. Biochem. 282, 29–38. 10.1006/abio.2000.4538

Lau, S., Rangarajan, R., Krüger-Genge, A., Braune, S., Küpper, J. H., Lendlein, A. and Jung, F. (2020). Age-related morphology and function of human arterial endothelial cells. Clin. Hemorheol. Microcirc. 74, 93–107. 10.3233/CH-199238

Lejeune, J., Levan, A., Böök, J. A., Chu, E. H. Y., Ford, C. E., Fraccaro, M., Harnden, D. G., Hsu, T. C., Hungerford, D. A., Jacobs, P. A. et al. (1960). A proposed standard system of nomenclature of human mitotic chromosomes. Lancet 275, 1063–1065 10.1016/S0140-6736(60)90948-X.

Lewis, J. P., Tanke, H. J., Raap, A. K., Beverstock, G. C. and Kluin-Nelemans, H. C. (1993). Somatic pairing of centromeres and short arms of chromosome 15 in the hematopoietic and lymphoid system. Hum. Genet. 92, 577–582. 10.1007/BF00420942

Lezhava, T. (2001). Chromosome and aging: genetic conception of aging. Biogerontology 2, 253–260. 10.1023/a:1013266411263

Lezhava, T. A. (1984). The activity of nucleolar organizer regions of human chromosomes in extreme old age. Gerontology 30, 94–99. 10.1159/000212614

Lima, M. F., Lisboa, M. O., Terceiro, L. E. L., Rangel-Pozzo, A. and Mai, S. (2022). Chromosome territories in hematological malignancies. Cells 11, 1368. 10.3390/cells11081368

Liu, Y., Bloom, S. I. and Donato, A. J. (2019). The role of senescence, telomere dysfunction and shelterin in vascular aging. Microcirculation 26, e12487. 10.1111/micc.12487

Liu, Z., Belmonte, J. C. I., Zhang, W., Qu, J. and Liu, G. H. (2022). Deciphering aging at three-dimensional genomic resolution. Cell Insight 1, 100034. 10.1016/j.cellin.2022.100034

López-Otín, C., Blasco, M. A., Partridge, L., Serrano, M. and Kroemer, G. (2013). The hallmarks of aging. Cell 153, 1194–1217. 10.1016/j.cell.2013.05.039

Loppin, B., Dubruille, R. and Horard, B. (2015). The intimate genetics of *Drosophila* fertilization. Open Biol. 5, 150076. 10.1098/rsob.150076

Maddox, A. S., Habermann, B., Desai, A. and Oegema, K. (2005). Distinct roles for two *C. elegans* anillins in the gonad and early embryo. Development 132, 2837–2848. 10.1242/dev.01828

Magidson, V., O’Connell, C. B., Lončarek, J., Paul, R., Mogilner, A. and Khodjakov, A. (2011). The spatial arrangement of chromosomes during prometaphase facilitates spindle assembly. Cell 146, 555–567. 10.1016/j.cell.2011.07.012

Martin, J. M., Kellett, J. M. and Kahn, J. (1980). Aneuploidy in cultured human lymphocytes: I. Age and sex differences. Age Ageing 9, 147–153. 10.1093/ageing/9.3.147

Mayer, W., Niveleau, A., Walter, J., Fundele, R. and Haaf, T. (2000). Demethylation of the zygotic paternal genome. Nature 403, 501–502.

Mayer, W., Fundele, R. and Haaf, T. (2000). Spatial separation of parental genomes during mouse interspecific (*Mus musculus* × *M. spretus*) spermiogenesis. Chromosome Res. 8, 555–558. 10.1023/a:1009227924235

Meaburn, K. J., Misteli, T. and Soutoglou, E. (2007). Spatial genome organization in the formation of chromosomal translocations. Semin. Cancer Biol. 17, 80–90. 10.1016/j.semcancer.2006.10.008

Misteli, T. (2007). Beyond the sequence: cellular organization of genome function. Cell 128, 787–800. 10.1016/j.cell.2007.01.028

Misteli, T. and Soutoglou, E. (2009). The emerging role of nuclear architecture in DNA repair and genome maintenance. Nat. Rev. Mol. Cell Biol. 10, 243–254. 10.1038/nrm2651

Miyai, T., Maruyama, Y., Osakabe, Y., Nejima, R., Miyata, K. and Amano, S. (2008). Karyotype changes in cultured human corneal endothelial cells. Mol. Vis. 14, 942–950.

Muller, W. A. and Gimbrone, M. A., Jr (1986). Plasmalemmal proteins of cultured vascular endothelial cells exhibit apical-basal polarity: analysis by surface-selective iodination. J. Cell Biol. 103, 2389–2402. 10.1083/jcb.103.6.2389

Nagele, R., Freeman, T., McMorrow, L. and Lee, H. Y. (1995). Precise spatial positioning of chromosomes during prometaphase: evidence for chromosomal order. Science 270, 1831–1835. 10.1126/science.270.5243.1831

Nagele, R. G., Freeman, T., Fazekas, J., Lee, K. M., Thomson, Z. and Lee, H. Y. (1998). Chromosome spatial order in human cells: evidence for early origin and faithful propagation. Chromosoma 107, 330–338. 10.1007/s004120050315

Nath, J., Tucker, J. D. and Hando, J. C. (1995). Y chromosome aneuploidy, micronuclei, kinetochores and aging in men. Chromosoma 103, 725–731. 10.1007/BF00344234

Naylor, R. M. and van Deursen, J. M. (2016). Aneuploidy in cancer and aging. Annu. Rev. Genet. 50, 45–66. 10.1146/annurev-genet-120215-035303

Nowinski, G. P., Van Dyke, D. L., Tilley, B. C., Jacobsen, G., Babu, V. R., Worsham, M. J., Wilson, G. N. and Weiss, L. (1990). The frequency of aneuploidy in cultured lymphocytes is correlated with age and gender but not with reproductive history. Am. J. Hum. Genet. 46, 1101–1111.

Odartchenko, N. and Keneklis, T. (1973). Localization of paternal DNA in interphase nuclei of mouse eggs during early cleavage. Nature 241, 528–529. 10.1038/241528a0

O’Toole, E., Greenan, G., Lange, K. I., Srayko, M. and Müller-Reichert, T. (2012). The role of γ-tubulin in centrosomal microtubule organization. PLoS One 7, e29795. 10.1371/journal.pone.0029795

Pierre, R. V. and Hoagland, H. C. (1972). Age-associated aneuploidy: loss of Y chromosome from human bone marrow cells with aging. Cancer 30, 889–894. 10.1002/1097-0142(197210)30:4%3C889::AID-CNCR2820300405%3E3.0.CO;2-1

Potapova, T. A., Unruh, J. R., Yu, Z., Rancati, G., Li, H., Stampfer, M. R. and Gerton, J. L. (2019). Superresolution microscopy reveals linkages between ribosomal DNA on heterologous chromosomes. J. Cell Biol. 218, 2492–2513. 10.1083/jcb.201810166

Reichmann, J., Nijmeijer, B., Hossain, M. J., Eguren, M., Schneider, I., Politi, A. Z., Roberti, M. J., Hufnagel, L., Hiiragi, T. and Ellenberg, J. (2018). Dual-spindle formation in zygotes keeps parental genomes apart in early mammalian embryos. Science 361, 189–193. 10.1126/science.aar7462

Richard, F., Muleris, M. and Dutrillaux, B. (1994). The frequency of micronuclei with X chromosome increases with age in human females. Mutat. Res. 316, 1–7. 10.1016/0921-8734(94)90002-7

Rippe, C., Blimline, M., Magerko, K. A., Lawson, B. R., LaRocca, T. J., Donato, A. J. and Seals, D. R. (2012). MicroRNA changes in human arterial endothelial cells with senescence: relation to apoptosis, eNOS and inflammation. Exp. Gerontol. 47, 45–51. 10.1016/j.exger.2011.10.004

Rodman, T. C., Flehinger, B. J. and Rohlf, F. J. (1980). Metaphase chromosome associations: Colcemid distorts the pattern. Cytogenet. Cell Genet. 27, 98–110. 10.1159/000131471

Rogers, M. J. (2021). Y chromosome copy number variation and its effects on fertility and other health factors: a review. Transl. Androl. Urol. 10, 1373–1382. 10.21037/tau.2020.04.06

Rohlf, F. J., Rodman, T. C. and Flehinger, B. J. (1980). The use of nonmetric multidimensional scaling for the analysis of chromosomal associations. Comput. Biomed. Res. 13, 19–35. 10.1016/0010-4809(80)90003-8

Sakurai, M. and Sandberg, A. A. (1976). The chromosomes and causation of human cancer and leukemia. XVIII. The missing Y in acute myeloblastic leukemia (AML) and Ph1-positive chronic myelocytic leukemia (CML). Cancer 38, 762–769. 10.1002/1097-0142(197210)30:4%3C889::AID-CNCR2820300405%3E3.0.CO;2-1

Schneider, I., de Ruijter-Villani, M., Hossain, M. J., Stout, T. A. E. and Ellenberg, J. (2021). Dual spindles assemble in bovine zygotes despite the presence of paternal centrosomes. J. Cell Biol. 220, e202010106. 10.1083/jcb.202010106

Shabanian, K., Shabanian, T., Karsai, G., Pontiggia, L., Paneni, F., Ruschitzka, F., Beer, J. H. and Saeedi Saravi, S. S. (2024). AQP1 differentially orchestrates endothelial cell senescence. Redox Biol. 76, 103317. 10.1016/j.redox.2024.103317

Srinivasan, D., Shisode, T., Shrinet, J. and Fraser, P. (2022). Chromosome organization through the cell cycle at a glance. J. Cell Sci. 135, jcs244004. 10.1242/jcs.244004

Stamatov, R., Uzunova, S., Kicheva, Y., Karaboeva, M., Blagoev, T. and Stoynov, S. (2025). Supra-second tracking and live-cell karyotyping reveal principles of mitotic chromosome dynamics. Nat. Cell Biol. 27, 654–667. 10.1038/s41556-025-01637-6

Stone, J. F. and Sandberg, A. A. (1995). Sex chromosome aneuploidy and aging. Mutat. Res. 338, 107–113. 10.1016/0921-8734(95)00016-y

Sun, H. B., Shen, J. and Yokota, H. (2000). Size-dependent positioning of human chromosomes in interphase nuclei. Biophys. J. 79, 184–190. 10.1016/S0006-3495(00)76282-5

Ting, K. K., Coleman, P., Zhao, Y., Vadas, M. A. and Gamble, J. R. (2021). The aging endothelium. Vasc. Biol. 3, R35–R47. 10.1530/VB-20-0013

Velez-Aguilera, G., Nkombo Nkoula, S., Ossareh-Nazari, B., Link, J., Paouneskou, D., Van Hove, L., Joly, N., Tavernier, N., Verbavatz, J. M., Jantsch, V. et al. (2020). PLK-1 promotes the merger of the parental genome into a single nucleus by triggering lamina disassembly. eLife 9, e59510. 10.7554/eLife.59510

Walczak, C. E., Cai, S. and Khodjakov, A. (2010). Mechanisms of chromosome behaviour during mitosis. Nat. Rev. Mol. Cell Biol. 11, 91–102. 10.1038/nrm2832

Weng, S., Stoner, S. A. and Zhang, D. E. (2016). Sex chromosome loss and the pseudoautosomal region genes in hematological malignancies. Oncotarget 7, 72356–72372. 10.18632/oncotarget.12050

Wiktor, A., Rybicki, B. A., Piao, Z. S., Shurafa, M., Barthel, B., Maeda, K. and Van Dyke, D. L. (2000). Clinical significance of Y chromosome loss in hematologic disease. Genes Chromosomes Cancer 27, 11–16.

Wood, J. G. and Helfand, S. L. (2013). Chromatin structure and transposable elements in organismal aging. Front. Genet. 4, 274. 10.3389/fgene.2013.00274

Wood, J. G., Hillenmeyer, S., Lawrence, C., Chang, C., Hosier, S., Lightfoot, W., Mukherjee, E., Jiang, N., Schorl, C., Brodsky, A. S. et al. (2010). Chromatin remodeling in the aging genome of *Drosophila*. Aging Cell 9, 971–978. 10.1111/j.1474-9726.2010.00624.x

Zhang, C., Zhang, X., Huang, L., Guan, Y., Huang, X., Tian, X. L., Zhang, L. and Tao, W. (2021). ATF3 drives senescence by reconstructing accessible chromatin profiles. Aging Cell 20, e13315. 10.1111/acel.13315

Zhang, L., Aviv, H., Gardner, J. P., Okuda, K., Patel, S., Kimura, M., Bardeguez, A. and Aviv, A. (2000). Loss of chromosome 13 in cultured human vascular endothelial cells. Exp. Cell Res. 260, 357–364. 10.1006/excr.2000.4997

Zhang, L. J., Shin, E. S., Yu, Z. X. and Li, S. B. (2007). Molecular genetic evidence of Y chromosome loss in male patients with hematological disorders. Chin. Med. J. 120, 2002–2005.

Zody, M. C., Garber, M., Adams, D. J., Sharpe, T., Harrow, J., Lupski, J. R., Nicholson, C., Searle, S. M., Wilming, L. and Young, S. K. et al. (2006). DNA sequence of human chromosome 17 and analysis of rearrangement in the human lineage. Nature 440, 1045–1049. 10.1038/nature04689

