## Supplemental Figures for "Higher frequency of homologous chromosome pairing in adult endothelial cells as compared to neonatal endothelial cells"

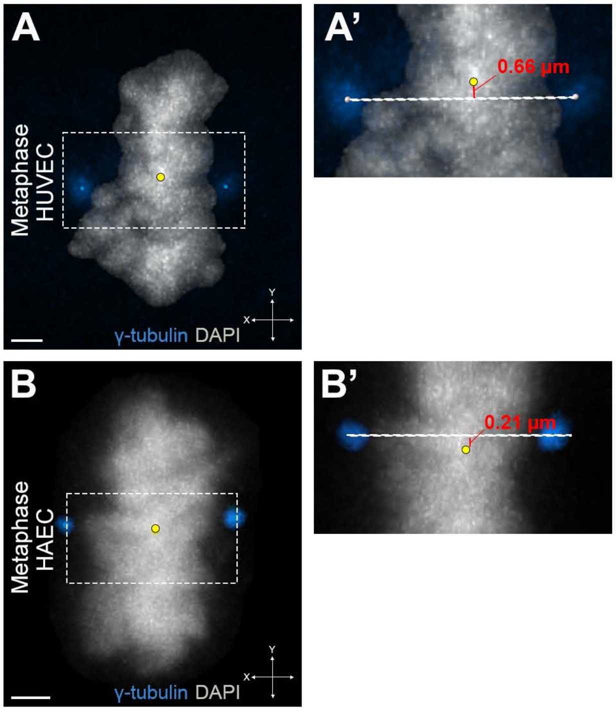


**Fig. S1. Characterizing the centrosome axis and center of 3D DNA surface in HUVECs and HAECs.**

**(A)** A metaphase HUVEC stained for γ-tubulin antibody (blue), and DNA (DAPI, grey). Yellow dot represents the center of volume of the DAPI stained chromosomes. Boxed region shows the x-, or centrosome axis. **(A')** Zoom in view of the boxed region in (A). Boundary zone (red line) is determined by measuring the distance between the chromosomal center of volume (yellow dot) and the centrosome axis (white dotted line). The distance between the centrosome axis (line connecting the centrosomes) and the chromosomal center of volume on average is ± 0.5 µm. Thus, a 1 µm width bounding box region of DAPI staining overlapping the chromosomal center of volume was determined as the "boundary zone" as previously described (Cai, Casas, Quintero Plancarte et al., 2025). **(B, B')** Same as (A, A'), but of HAECs. The average distance between the centrosome axis and the chromosomal center of volume is ± 0.45 µm. Therefore the boundary region is 0.90 µm for HAECs. Note: For chromosome painting analysis, if the center of volume of an individual homolog was ± 0.5 µm for HUVECs and ± 0.45 for HAECs from the chromosomal center of volume along the x-axis, these cells were not used in our analysis as they were uninterpretable. For instance, out of 66 total cells, 6 cells exhibited homologous chromosome 1 in this boundary region, rendering the data uninterpretable (n=6/66 cells for chromosome 1). Scale bar: 2 µm.

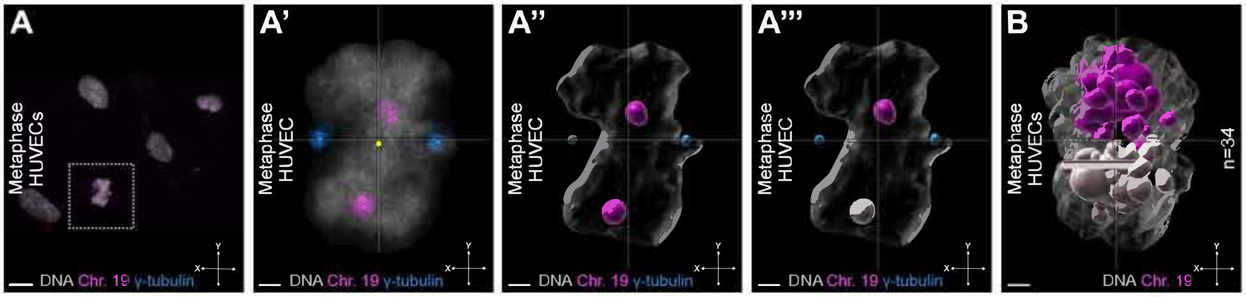


**Fig. S2. Generation of a 3D overlay and analysis.**
**(A)** Top view of the confocal staining at 63x magnification of HUVECs at interphase and metaphase stains for DNA (grey), γ-tubulin (blue), and chromosome 19 (magenta). Boxed region shows a mitotic cell. **(A')** As in (A), but a zoom in of the boxed region in (A) showing segregation of homologous chromosome 19. Center of the DAPI stained chromosomes (yellow dot) and centrosomes (blue dot) are aligned along the centrosome axis. Center of volume for each chromosome 19 is also shown (magenta dot). **(A")** A 3D reconstruction of optical confocal sections in (A'). **(A'")** To differentiate between each homologous chromosome, the one homolog that is closest to the x-, or centrosome, axis is designated in color (magenta), while its respective homolog is in grey. **(B)** Generation of a 3D overlay to determine if each pair of homologous chromosomes are spatially segregated from one another. Individual mitotic cells overlaid using the centrosome axis as previously described (Hua and Mikawa, 2018b). For each metaphase cell, we determined if they were spatially segregated by using the centrosome axis. If the homologous chromosomes are located on opposite sides, it indicates spatial segregation of the homologs. In contrast, if both homologous chromosomes are positioned on the same side along the centrosome axis, it signifies a loss of segregation. Scale bars: 10, 2 μm.

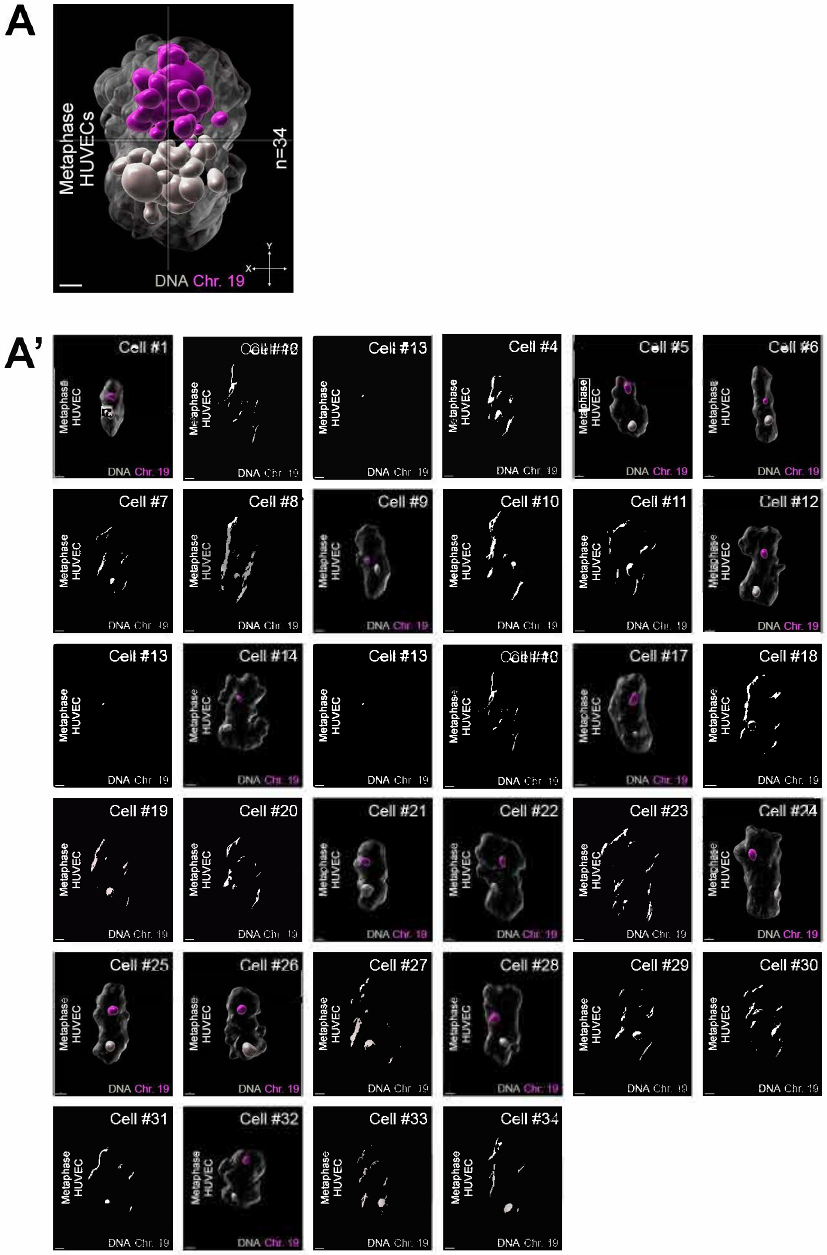


**Fig. S3 Composition of a 3D overlay for HUVECs and HAECs.**
**(A)** 3D overlay of HUVECs at metaphase as described in Fig. S2B (n=34 cells). **(A’)** Composition of individual HUVECs as described in Fig. S2A’’-A’’’). Note: Cell number is not equivalent to the cell number in the relative position graph in Fig. 1. **(B, B’)** As in (A, A’), but of HAECs (n=30 cells). Scale bar: 2 µm.

B

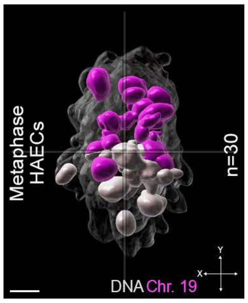


B’

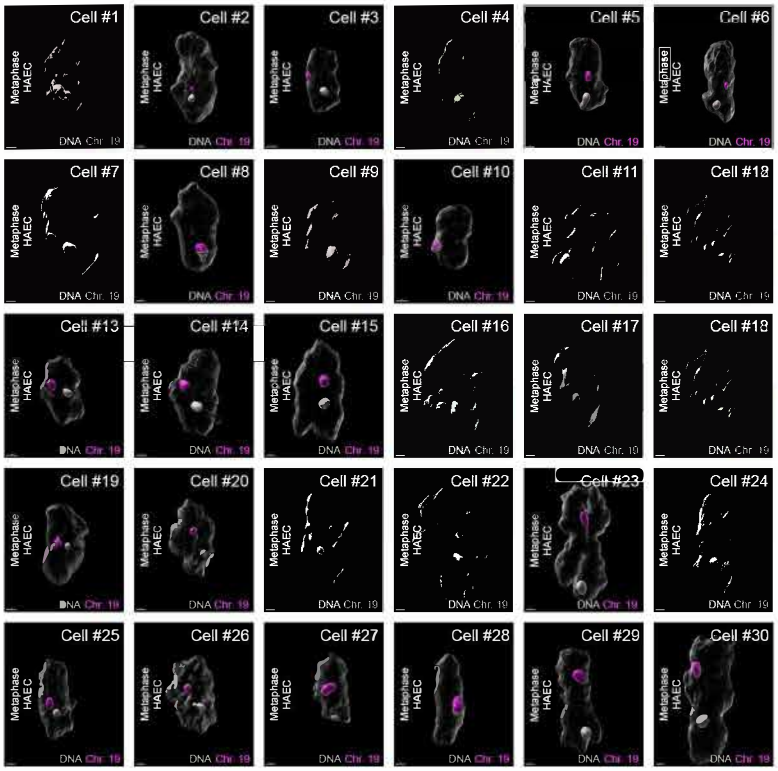


**Fig. S3 Composition of a 3D overlay for HUVECs and HAECs.**
**(A)** 3D overlay of HUVECs at metaphase as described in Fig. S2B (n=34 cells). **(A’)** Composition of individual HUVECs as described in Fig. S2A’’-A’’’). Note: Cell number is not equivalent to the cell number in the relative position graph in Fig. 1. **(B, B’)** As in (A, A’), but of HAECs (n=30 cells). Scale bar: 2 µm.

Table S1. Analysis of various chromosomes for pooled HUVECs and individual HAEC patients

| Human Umbilical Vein Endothelial Cells (HUVECs) | Chr. 1 | Chr. 4 | Chr. 13 | Chr. 15 | Chr. 17 | Chr. 19 | Chr. 21 | Chr. 22 | Chr. X | Chr. Y |
|---|---|---|---|---|---|---|---|---|---|---|
| Total neonatal samples (20 patients) | 31 | 28 | 33 | 41 | 17 | 34 | 32 | 22 | 19 | 16 |
| **Human Aortic Endothelial Cells (HAECs)** | **Chr. 1** | **Chr. 4** | **Chr. 13** | **Chr. 15** | **Chr. 17** | **Chr. 19** | **Chr. 21** | **Chr. 22** | **Chr. X** | **Chr. Y** |
| Total adult samples | 60 | 49 | 47 | 48 | 83 | 30 | 51 | 99 | 132 | 44 |
| Patient #1: 21 Year Old Male | 7 | 3 | N/A | N/A | N/A | N/A | N/A | 6 | N/A | 16 |
| Patient #2: 36 Year Old Female | 18 | 10 | 23 | 25 | 46 | 6 | 25 | 1 | 37 | N/A |
| Patient #3: 45 Year Old Female | N/A | N/A | N/A | N/A | N/A | N/A | N/A | 48 | 46 | N/A |
| Patient #4: 50 Year Old Male | 6 | 1 | 12 | N/A | N/A | 4 | 14 | 8 | N/A | 9 |
| Patient #5: 50 Year Old Female | 27 | 30 | N/A | 15 | 4 | N/A | N/A | 21 | 38 | N/A |
| Patient #6: 50 Year Old Female | 2 | 5 | 4 | N/A | 12 | 13 | 3 | 14 | 11 | N/A |
| Patient #7: 68 Year Old Male | N/A | N/A | 8 | 8 | 21 | 7 | 9 | 1 | N/A | 19 |

**Table S1. Analysis of various chromosomes for pooled HUVECs and individual HAEC patients.**

Number of cells collected and analyzed from pooled HUVECs (n=20 patients) and individual HAECs (n=7 individual patients) for chromosomes 1 (magenta), 4 (green), 13 (red), 15 (green), 17 (cyan), 19 (magenta), 21 (magenta), 22 (yellow), X (red), and Y (blue).

Table S2. Frequency of loss of spatial segregation of various chromosomes for pooled HUVECs and individual HAEC patients

| Human Umbilical Vein Endothelial Cells (HUVECs) | Chr. 1 | Chr. 4 | Chr. 13 | Chr. 15 | Chr. 17 | Chr. 19 | Chr. 21 | Chr. 22 | Chr. X | Chr. Y |
|---|---|---|---|---|---|---|---|---|---|---|
| Total neonatal samples (20 patients) | 6.45% | 7.14% | 0.00% | 19.5% | 0.00% | 2.94% | 15.6% | 18.2% | 0.00% | 18.8% |
| **Human Aortic Endothelial Cells (HAECs)** | **Chr. 1** | **Chr. 4** | **Chr. 13** | **Chr. 15** | **Chr. 17** | **Chr. 19** | **Chr. 21** | **Chr. 22** | **Chr. X** | **Chr. Y** |
| Total adult samples | 20.0% | 30.6% | 53.2% | 60.4% | 47.0% | 43.3% | 41.2% | 51.5% | 37.9% | 50.0% |
| Patient #1: 21 Year Old Male | 42.9% | 33.3% | N/A | N/A | N/A | N/A | N/A | 33.3% | N/A | 50.0% |
| Patient #2: 36 Year Old Female | 22.2% | 40.0% | 43.5% | 60.0% | 41.3% | 66.7% | 52.0% | 100% | 21.6% | N/A |
| Patient #3: 45 Year Old Female | N/A | N/A | N/A | N/A | N/A | N/A | N/A | 56.3% | 67.4% | N/A |
| Patient #4: 50 Year Old Male | 16.7% | 100% | 41.7% | N/A | N/A | 0.00% | 28.6% | 50.0% | N/A | 33.3% |
| Patient #5: 50 Year Old Female | 11.1% | 26.7% | N/A | 66.7% | 0.00% | N/A | N/A | 38.1% | 21.1% | N/A |
| Patient #6: 50 Year Old Female | 50.0% | 20.0% | 75.0% | N/A | 58.3% | 46.2% | 33.3% | 64.3% | 27.3% | N/A |
| Patient #7: 68 Year Old Male | N/A | N/A | 87.5% | 50.0% | 61.9% | 42.9% | 33.3% | 0.00% | N/A | 57.9% |

**Table S2. Frequency of loss of spatial segregation of various chromosomes for pooled HUVECs and individual HAEC patients.**

Percentage of HUVECs (n=20 patients) and individual HAECs (n=7 individual patients) that have lost spatial segregation for chromosomes 1 (magenta), 4 (green), 13 (red), 15 (green), 17 (cyan), 19 (magenta), 21 (magenta), 22 (yellow), X (red), and Y (blue).

Table S3. Summary of the mean minimal distance of homologous pairs of HUVECs and HAECs

| Human Umbilical Vein Endothelial Cells (HUVECs) | Chr. 1 | Chr. 4 | Chr. 13 | Chr. 15 | Chr. 17 | Chr. 19 | Chr. 21 | Chr. 22 | Chr. X | Chr. Y |
|---|---|---|---|---|---|---|---|---|---|---|
| Average Distance (µm) | 3.25 | 3.67 | 3.19 | 1.88 | 3.03 | 2.98 | 2.21 | 1.50 | 3.79 | 2.42 |
| Standard Deviation | 1.45 | 1.48 | 1.27 | 1.72 | 0.99 | 1.70 | 1.60 | 1.09 | 1.08 | 1.06 |
| **Human Aortic Endothelial Cells (HAECs)** | **Chr. 1** | **Chr. 4** | **Chr. 13** | **Chr. 15** | **Chr. 17** | **Chr. 19** | **Chr. 21** | **Chr. 22** | **Chr. X** | **Chr. Y** |
| Average Distance (µm) | 3.45 | 3.38 | 2.06 | 1.67 | 1.68 | 1.99 | 1.81 | 1.28 | 2.79 | 2.04 |
| Standard Deviation | 2.30 | 2.88 | 1.66 | 1.85 | 1.40 | 1.89 | 1.35 | 1.13 | 2.57 | 1.44 |

**Table S3. Summary of the mean minimal distance of homologous pairs of HUVECs and HAECs.**

The mean and standard deviation of the minimal distance between homologous chromosomes of HUVECs and HAECs for chromosomes 1 (magenta), 4 (green), 13 (red), 15 (green), 17 (cyan), 19 (magenta), 21 (magenta), 22 (yellow), X (red), and Y (blue).

Table S4. Summary of the mean angular orientation of homologous pairs of HUVECs and HAECs

| Human Umbilical Vein Endothelial Cells (HUVECs) | Chr. 1 | Chr. 4 | Chr. 13 | Chr. 15 | Chr. 17 | Chr. 19 | Chr. 21 | Chr. 22 | Chr. X | Chr. Y |
|---|---|---|---|---|---|---|---|---|---|---|
| Average Vector Angle | 48.2° | 43.9° | 39.7° | 27.5° | 34.5° | 5.61° | 15.7° | 4.56° | 26.5° | 29.5° |
| Standard Deviation | 19.3° | 30.8° | 29.4° | 27.7° | 21.9° | 4.27° | 20.3° | 2.73° | 26.7° | 25.5° |
| **Human Aortic Endothelial Cells (HAECs)** | **Chr. 1** | **Chr. 4** | **Chr. 13** | **Chr. 15** | **Chr. 17** | **Chr. 19** | **Chr. 21** | **Chr. 22** | **Chr. X** | **Chr. Y** |
| Average Vector Angle | 45.8° | 48.6° | 20.8° | 26.6° | 27.2° | 5.43° | 8.78° | 12.6° | 35.2° | 27.2° |
| Standard Deviation | 32.3° | 30.1° | 26.6° | 29.3° | 27.5° | 4.53° | 12.5° | 20.0° | 26.7° | 24.6° |

**Table S4. Summary of the mean angular orientation of homologous pairs of HUVECs and HAECs.**

The mean and standard deviation of the angular orientation (vector angle) between homologous chromosomes of HUVECs and HAECs for chromosomes 1 (magenta), 4 (green), 13 (red), 15 (green), 17 (cyan), 19 (magenta), 21 (magenta), 22 (yellow), X (red), and Y (blue).

**Movie 1. Homologous segregation is lost in acrocentric chromosome 13 in adult HAECs as compared to neonatal HUVECs.**

Video of Whole Chromosome Painting of chromosome 13 and 3D overlays for HUVECs (top panels), and HAECs (bottom panels) rotating 360º counter-clockwise along the y-axis staining for chromosome 13 (red), and DNA (SYTOX, grey) for whole chromosome painting, and chromosome 13 surfaces (red/grey) for overlays. Scale bar: 3 μm.

### References

Cai, P., Casas, C. J., Quintero Plancarte, G., Mikawa, T. and Hua, L. L. (2025). Ipsilateral restriction of chromosome movement along a centrosome, and apical-basal axis during the cell cycle. Chromosome Res. 33, 1-21. https://doi.org/10.1007/s10577-024-09760-0

Hua, L. L. and Mikawa, T. (2018). Mitotic antipairing of homologous and sex chromosomes via spatial restriction of two haploid sets. Proc. Natl. Acad. Sci. USA 115, E12235-E12244. https://doi.org/10.1073/pnas.1809583115
